# Expanding the GUSome: Structure-guided identification and characterization of gut microbial β-glucuronidases

**DOI:** 10.64898/2026.06.20.733316

**Authors:** Tarushi, Chinmaya V Badgujar, Subhash C Bihani

## Abstract

The gut microbiome-encoded β-glucuronidase (GUS) enzymes have a significant effect on human physiology through their deglucuronidation activity on endogenous and exogenous glucuronides. GUS activity also significantly influences the pharmacokinetics, efficacy and toxicity of various drugs including chemotherapeutic drugs. Given their crucial role in drug metabolism, GUS enzymes have emerged as promising targets for therapeutic intervention. Here, we have identified and characterized 79 unique GUS enzymes through a structure-guided approach. Structural modelling of these GUS enzymes revealed a conserved core and active-site residues with significant variations in the number and nature of the C-terminal domains. A new classification system based on the number and type of additional C-terminal domains is presented for the GUS proteins. Further, GUS enzymes have been categorized into different loop categories linked to their substrate preferences. The relationship between domain architecture and loop-type is explored by sequence similarity network analysis. We could successfully express, purify and validate GUS processing capability of a panel of identified GUS proteins. The nature of oligomer organization has been deciphered by SEC and DLS studies. Further, we have identified additional GUS enzymes capable of processing SN-38G, glucuronidated form of anticancer drug, irinotecan. These newly identified GUS enzymes will offer valuable insights into gut microbial GUS diversity and their role in understanding the population-specific drug-induced adverse effects on human health.

## Introduction

The human gastrointestinal tract serves as a host to a complex community of microbes known as the gut microbiota, which collectively codes for a rich diversity of enzymes including carbohydrate-active enzymes (CAZymes) (Kaoutari et al., 2013; Wardman et al., 2022). Among them, “GUSome”, the collection of β-glucuronidase enzymes, holds a unique position in both microbial and host physiology. These enzymes belonging to the glycoside hydrolase 2 (GH2) family, catalyse the removal of glucuronic acid from a wide variety of host-derived and exogenous compounds including complex structural glycosaminoglycans, dietary carbohydrates, endogenous hormones, and xenobiotic glucuronides (Pellock & Redinbo, 2017; Wallace et al., 2010). Through their activity, GUS enzymes dictate the bioavailability, half-life, and toxicity of numerous biologically active compounds.

One of the well-characterized roles of GUS in the context of human pathophysiology is reactivation of drugs and toxic xenobiotics. Glucuronidation by human uridine diphosphate glucuronosyltransferases (UGTs) is a major detoxification pathway that promotes excretion of drugs and toxic xenobiotics (G. F. Dutton, 1966; G. J. Dutton, 2019). Deglucuronidation by the microbial GUS in the gut regenerates parent aglycones, which can be reabsorbed through the intestinal mucosa, entering the portal circulation and returning to the liver in a process known as the enterohepatic recirculation (Vítek & Carey, 2003). This process extends the biological half-life of compounds, which can be beneficial for some nutrients but highly problematic for other metabolites leading to local toxicity. A prominent and clinically significant example of GUS-mediated toxicity involves irinotecan, a first-line chemotherapeutic agent used for colorectal and pancreatic cancers. In the liver, the active drug metabolite SN-38 is detoxified by Human UGTs to form the inactive metabolite SN-38 glucuronide (SN-38G), which is subsequently excreted into the gastrointestinal tract. Gut microbial GUS enzymes hydrolyze SN-38G to produce SN-38 directly in the intestinal lumen which causes epithelial damage and severe, dose-limiting diarrhea (Takasuna et al., 1996; Wallace et al., 2010; Roberts et al., 2013). Involvement of GUS activity has also been established in the regulation of non-steroidal anti-inflammatory drugs (NSAIDs), e.g., diclofenac (Zhong et al., 2016), hormones, e.g., estrogen (Ervin, Li, et al., 2019) and immunosuppressant, e.g., mycophenolate mofetil (MMF) (Spanogiannopoulos et al., 2016; Xia et al., 2024). Moreover, in some studies, GUS activity has been linked to increased risk of cancer development, e.g., breast cancer, tobacco-induced cancers (Ervin, Li, et al., 2019; Greten & Arkan, 2024). GUS enzymes are also involved in the metabolism of dietary polyphenols and flavonoids,in the human gut influencing the nutrient bioavailability (Rowland et al., 2018; Selma et al., 2009). Their deglucuronidation activity has been exploited for the production of bioactive metabolites such as glycyrrhetinic acid monoglucuronide from its diglucuronide conjugate, i.e., glycyrrhizin. The GUS enzyme from *Escherichia coli* has also been extensively used as a reporter gene in gene fusion studies in plants (Jefferson et al., 1987).

Due to their significant role in host physiology, potential therapeutic intervention and biotechnological applications, it is important to understand GUSome diversity and establish a clear structure-function relationship. The structure of microbial GUS, first determined for *E. coli* GUS (EcGUS), showed a conserved GH2 core consisting of three domains: an N-terminal jelly-roll domain, an immunoglobulin-like (Ig-like) domain, and a C-terminal TIM-barrel domain that houses the active site (Wallace et al., 2010). The active site is defined by the presence of seven key conserved active site residues termed “GUS rubric” (Pollet et al., 2017). The GH2 core is essentially conserved among the bacterial GUS enzymes with structural and functional diversity arising from the active site loops and the presence of additional C-terminal domains. Bacterial GUS enzymes have been classified into eight distinct structural categories based on the architecture of active site loops (Walker et al., 2022). These loops have been shown to dictate the substrate specificity of GUS enzymes, specifically with respect to small-molecule drug glucuronides. Further, C-terminal domains have also been shown to influence substrate processing; however, their role is less characterized.

To further expand the structural-functional landscape of GUSome, we conducted a comprehensive bioinformatics analysis of a metagenomic dataset representing the Indian gut microbiome and identified 79 unique gut microbial GUS enzymes. Based on extensive domain analysis, a new classification system based on domain architecture is proposed. Detailed examination of different active-site loops is performed and their association with domain architecture is evaluated. To validate our in-silico findings, representative GUS proteins were expressed, purified and characterized biochemically to validate their β-glucuronidase activity. Crucially, some of these newly characterized GUS enzymes were found to actively cleave chemotherapeutic metabolite SN-38G, enhancing our understanding of the specific microbial enzymes driving irinotecan toxicity and expanding the catalogue of functionally characterized GUS enzymes relevant to human health.

### Results and discussions Identification of GUS enzymes

β-glucuronidases specialize in the removal of terminal β-D-glucuronic acid from diverse substrates and show high specificity towards glucuronic acid while accepting wide variations in the aglycone moiety. A set of seven conserved active site residues, i.e., Asn412, Glu413, Tyr468, Glu504, Asn566, Lys568, and Gly569 (corresponding to EcGUS), collectively known as ‘GUS rubric’, are essential for this specific glucuronidase activity (Figure 1A) (Wallace et al., 2010). The presence of a complete GUS rubric, therefore, defines a functional β-glucuronidase and differentiates it from other glycoside hydrolases. Glu413 and Glu504 are catalytic residues while the specificity for the glucuronide over other sugars is provided by the residues Asn566, Lys568, and Gly569, popularly known as “N*KG motif” (Wallace et al., 2010; Wallace et al., 2015). This is achieved by specific interactions of residues Asn566 and Lys568 with the carboxylate moiety present at the C6 position of glucuronides (Figure 1A). Additionally, Tyr468 provides specificity towards β-linked substrates through steric occlusion of ɑ-linked substrates (Wallace et al., 2015). Recently, an “extended GUS rubric” has been proposed where Tyr468 is replaced by a tryptophan residue while maintaining the required interactions (Pellock et al., 2018). In this study, we have used the presence of this extended GUS rubric as the defining criterion for the identification of true GUS enzymes.

**Figure 1:**
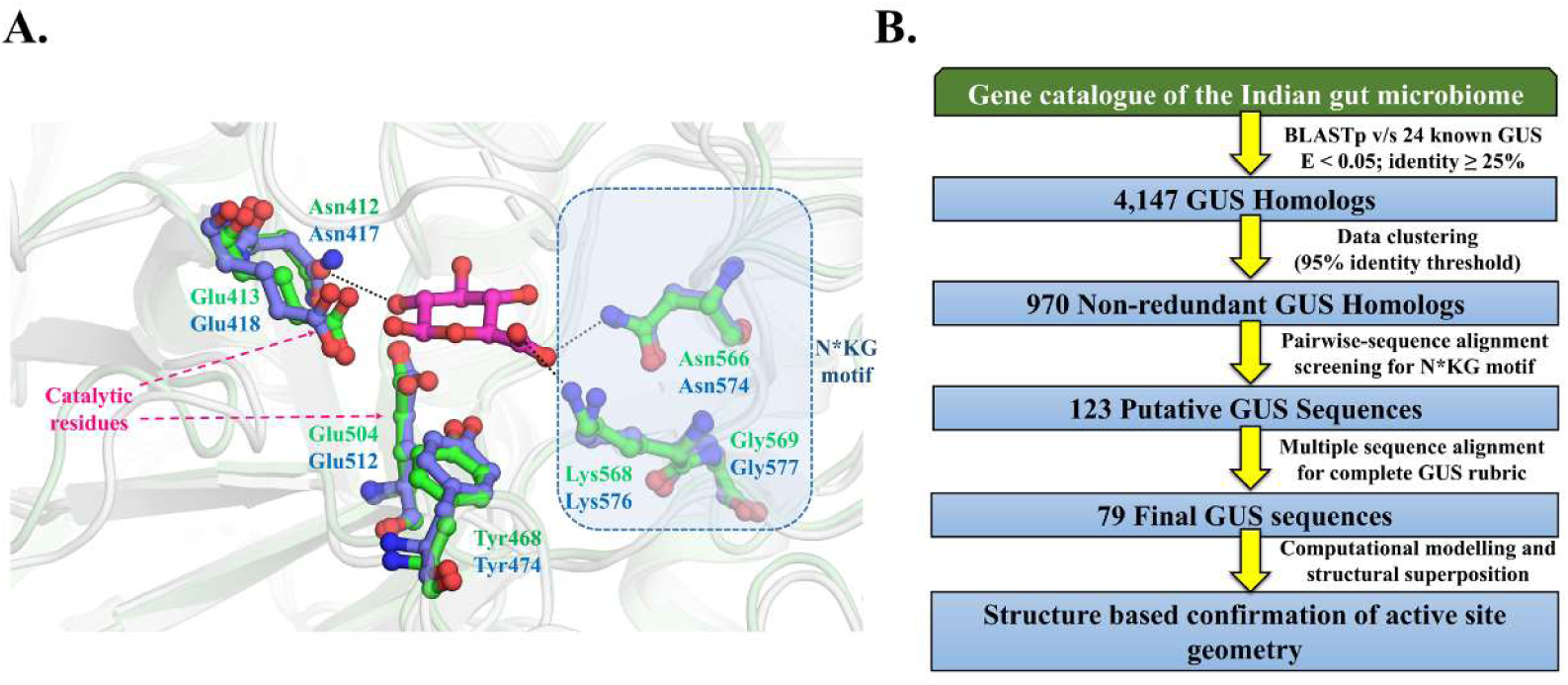
GUS rubric residues and identification of ‘true’ GUS enzymes: (A) Active site of EcGUS (PDB ID: 6LEM) depicting GUS rubric residues in ball and stick mode (*green color*). The N*KG motif (Asn566, Lys568, and Gly569) is enclosed in a blue frame. RgGUS (PDB ID: 6JZ5) active site residues (Asn417, Glu418, Tyr474, Glu512, Asn574, Lys576, and Gly577, *slate*) in complex with glucuronide moiety are also shown to highlight the interactions with the glucuronide moiety (*magenta*). (B) Bioinformatics workflow depicting the steps for the identification of gut microbial GUS homologs. The number of sequences remaining after each step is indicated in the boxes.

A representative gene catalogue of the Indian gut microbiome was chosen to identify GUS enzymes prevalent in Indian population (Dhakan et al., 2019). GUS homologs were identified through a multi-step bioinformatics pipeline (Figure 1B). First, a BLASTp search was performed against the gene catalogue using protein sequences of 24 structurally characterized GUS as queries. This step identified 4,147 sequences and further clustering at 95% sequence identity resulted in 970 non-redundant GUS homologs. To identify ‘true’ GUS enzymes, these 970 sequences were manually screened for the presence of the N*KG motif in a pairwise alignment with the known GUS proteins, resulting in 123 sequences. Multiple sequence alignment (MSA) using Clustal Omega was conducted to verify the presence of the complete 7-residue GUS rubric (Sievers et al., 2011). This step ultimately yielded 79 putative GUS sequences with a complete GUS rubric and a properly aligned N*KG motif (Supplementary Figure 1). Further, computational structural models of all the 79 GUS enzymes were prepared and the presence of a properly organized GUS active site was confirmed in every model through structural superposition and visual inspection.

### Taxonomic profiling

Gut microbial GUS enzymes have been shown to be distributed among diverse bacterial taxa including Bacteroidota, Firmicutes, Proteobacteria, Verrucomicrobiota and Actinobacteriota (Pollet et al., 2017). Similarly, in the present study also, maximum GUS sequences belonged to phylum Bacteroidota (53%) followed by Firmicutes (40%) (Supplementary Figure 2). GUS enzymes from several commensal taxa prevalent in the Indian gut ecosystem including *Bacteroides uniformis, Bifidobacterium adolescentis, Faecalibacterium prausnitzii, Prevotella sp., Segatella copri and Escherichia coli*, were identified. *Segatella copri* is one of the most prevalent gut symbionts in the Indian microbiome. In the present study, five GUS sequences belonging to the *Segatella copri* were identified. These observations provide insights into the taxonomic diversity of GUS enzymes within the Indian gut microbiome and highlight the need for further research into the population-specific role of these enzymes in nutrition and drug metabolism.

### Domain architecture and conserved GH2 core

Gut microbial GUS enzymes have a conserved GH2 core consisting of a TIM-barrel scaffold with adaptations including topological alterations, loop insertions, and fusion of additional domains. The catalytic TIM-barrel adopts a canonical (β/α)_8_ fold with the β/α loops or front loops forming the active-site wall and the α/β loops likely contribute to the structural stability (Sterner & Höcker, 2005) (Figure 2A). The TIM-barrel domain is extended at the N-terminus by two β-sandwich domains modulating the size and shape of the active site. These domains are similar to the domains described for GH2 β-galactosidases (Juers et al., 2000; Talens-Perales et al., 2016). The first domain (residues 1–180 in EcGUS) adopts a jelly-roll fold resembling the sugar-binding domains of other GH2 family CAZymes (Figure 2B) (Wallace et al., 2010). Region 181-273 in EcGUS folds into an immunoglobulin-like (Ig-like) domain and is likely to serve as a structural bridge between the jelly-roll and TIM-barrel domain (Figure 2B). While this Ig-like domain is highly conserved in GUS enzymes, it is replaced by an uncharacterized domain in some GH2 β-galactosidases (Talens-Perales et al., 2016).

**Figure 2:**
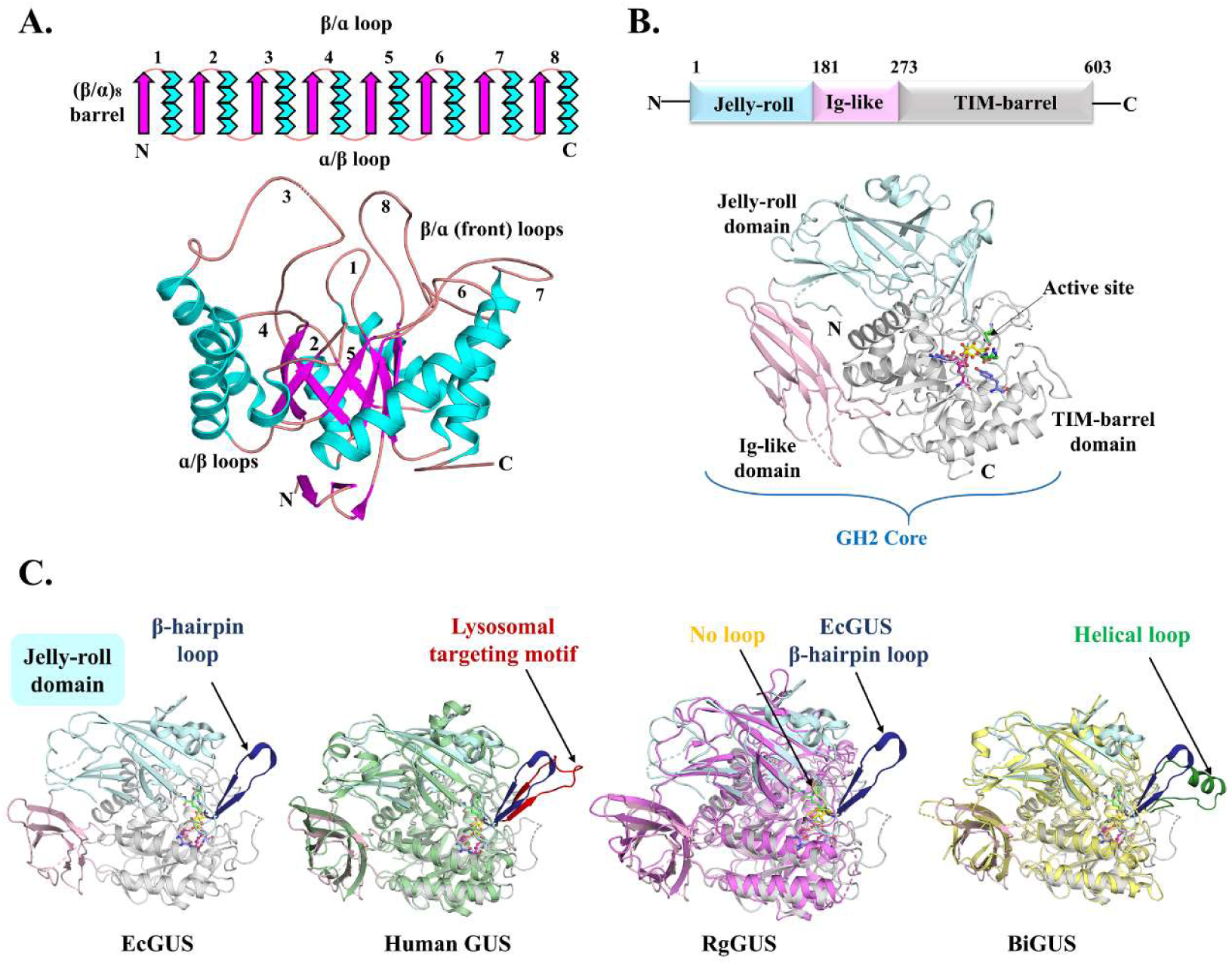
Domain architecture of GUS enzymes: (A) Topological diagram and cartoon representation of TIM-barrel domain of EcGUS showing (β/ɑ)_8_-barrel architecture and connecting loops. (B) The conserved glycoside hydrolase 2 (GH2) core of EcGUS includes N-terminal Jelly-roll domain (*pale cyan*), Ig-like domain (*light pink*) and C-terminal TIM-barrel domain (*grey*). (C) Presence of β-hairpin loop insertion in jelly-roll domain of EcGUS (*dark blue*), Human GUS (hGUS; PDB ID: 3HN3; *red*), no insertion in *Ruminococcus* GUS (RgGUS; PDB ID: 6MVG, *yellow*), and helical insertion in *Bifidobacterium dentium* GUS (BiGUS; PDB ID: 6LD6; *green*). EcGUS with β-hairpin loop insertion in *dark blue* is also shown for comparison.

A less explored structural feature of GUS enzymes is a β-hairpin loop insertion just before the last β-strand of the jelly-roll domain (Dashnyam et al., 2018). In human GUS, this β-hairpin loop serves as a lysosomal targeting motif, essential for enzyme localization (Figure 2C) (Naz et al., 2013). EcGUS (PDB ID: 6LEM), RgGUS (PDB ID: 5Z18) and CpGUS (PDB ID: 4JKM) possess similar but shorter loop with elongated β-strands. However, majority of structurally characterized GUS enzymes like Rg3GUS (PDB ID: 6MVG) and those from the present study lack this feature entirely (Figure 2C). Interestingly, some enzymes such as BiGUS (*Bifidobacterium dentium*; PDB ID: 6LD6) and Fp2-NL GUS (*Faecalibacterium prausnitzii*; PDB ID: 6U7I) and few GUS enzymes from the present study possess a larger loop with an α-helix, suggesting an alternative structural motif (Figure 2C). By analogy with human GUS, this loop insertion in microbial GUS may influence enzyme localization or due to its proximity to the active site potentially affect enzymatic activity or substrate specificity. In BiGUS (PDB ID: 6LD6) and EeGUS (PDB ID: 6BJW), residues from this loop region are suitably positioned near the active site to interact with the aglycone moiety of the substrate (Supplementary Figure 3) (Dashnyam et al., 2018). Deletion of this region resulted into significant loss of catalytic activity in EeGUS supporting its functional relevance (Biernat et al., 2019). However, the exact role of this insertion remains unclear and warrants further biochemical and localization studies.

### Diversity of C-terminal domains and a new classification of GUS enzymes

Overall, the GH2 core is well conserved among the structurally characterized GUS enzymes, with most variations localized primarily in the loop regions. However, several larger GUS enzymes have acquired one or more additional domains at the C-terminus. Using dbCAN3, Interpro, and structural analysis of computational models, we identified five distinct domain architectures among 24 structurally characterized GUS and 79 GUS enzymes from the present study. Based on the topological diversity of C-terminal domains, we propose a new classification of microbial GUS enzymes into five categories (Table 1, Figure 3).

**Figure 3:**
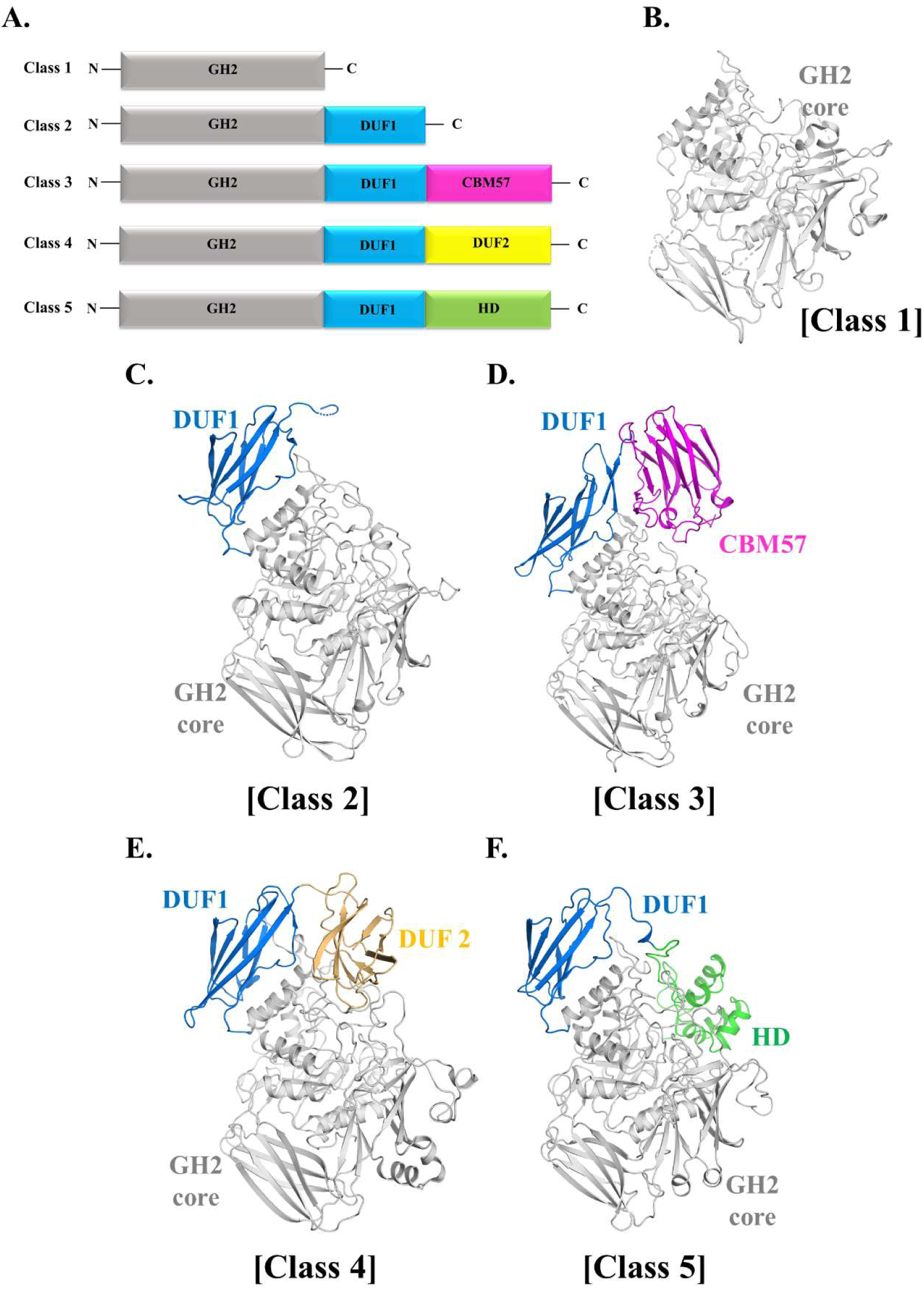
Classification of GUS enzymes based on C-terminal domains: (A) Five different classes of the gut microbial GUS enzymes based on the C-terminal domain diversity. (B) Class 1 with only GH2 core domain (EcGUS, *grey*) (C) Class 2 with an additional DUF1 domain (BfGUS, PDB ID: 3CMG, *blue*) (D) Class 3 with DUF1 and CBM57 domains (Bu2GUS, PDB ID: 5UJ6, *magenta*) (E) Class 4 with DUF1 and DUF2 domains (PmGUS, PDB ID: 6D7J, *golden*) (F) Class 5 with DUF1 and helical domains (*F. prausnitzii* GUS, HBV_6285 (model), *green*). GH2: Glycoside hydrolase 2, DUF: Domain of unknown function, CBM57: Carbohydrate binding module 57; HD: Helical domain.

**Table 1:** Classification of GUS enzymes in five classes based on domain architecture.

| **Classes** | **Number of additional C-terminal domains** | **First domain** | **Second domain** | **Representative PDB ID** |
| --- | --- | --- | --- | --- |
| Class 1: Minimal Core | 0 | NA | NA | 7PR6 |
| Class 2: Single Extension- DUF1 | 1 | DUF4982 | NA | 3CMG |
| Class 3: Double Extension - Malectin-like domain | 2 | DUF4982 | CBM57 | 5UJ6 |
| Class 4: Double Extension - DUF2 | 2 | DUF4982 | DUF2 | 6D7J |
| Class 5: Double Extension - helical domain | 2 | DUF4982 | Helical | NA |

Class 1 (Minimal core), represented by EcGUS, lacks any C-terminal extensions and includes 15 of 24 solved structures and 20 GUS from the present study (Figure 3B). Remaining GUS enzymes have one or more domains at the C-terminus. Interestingly, the first C-terminal domain in all the remaining GUS enzymes is identical (DUF4982), while there are variations in the type of second C-terminal domain. Based on these variations, the following classes have been identified: Class 2 (single extension-DUF1) is characterized by a single DUF4982 domain termed as DUF1 (Figure 3C) (Pellock et al., 2018, 2019). Five out of 24 structures, including BfGUS (PDB ID: 3CMG) and 30 GUS enzymes from present study belong to this class. Class 3 (double extension-CBM57) features a DUF1 domain followed by a carbohydrate-binding module 57 (CBM57) or malectin-like domain observed in Bu2GUS (PDB ID: 5UJ6) and BdGUS (PDB ID: 6ED1) (Figure 3D) (Pellock et al., 2018). Thirteen GUS enzymes from the present study could be placed in this category. Class 4 (double extension-DUF2) harbours a distinct DUF2 domain in place of CBM57 (Figure 3E) (Little et al., 2018). PmGUS (PDB ID: 6D7J) and BuGUS3 (PDB ID: 6D1P) typify this configuration and four GUS enzymes from the present study have this configuration. While CBM57 domain in BuGUS2 is positioned closer to the active site of the enzyme, the DUF2 domain in PmGUS is oriented away from it (Little et al., 2018; Pellock et al., 2018).

Analysis of 79 GUS from the present study also revealed a previously undescribed category of GUS enzymes containing predominantly helical C-terminal domains as compared to the β-sheet rich domains observed so far. There is substantial topological diversity among these helical domains (Supplementary figure 4). We have classified them as class 5 (double extension - helical domain) (Figure 3F). Foldseek analysis indicated that the smaller (∼100–120 residue) helical domains found in some (12) of the GUS enzymes (such as HBG_1439, HBV_6285, and HBI_1240, etc.) resemble the C-terminal domain observed in GH3 β-glucosidase from *Clostridium thermocellum* (PDB ID: 7MS2) (Supplementary Figure 5A). The function of this small domain is not yet elucidated. However, the domain is located close to the active site and in HBG_1439, a loop from this domain extends towards the active site suggesting a role in substrate processing (Supplementary Figure 5B). In contrast, the larger (∼200 residue) helical domain identified in *Bifidobacterium adolescentis* GUS (HCQ_7399) share very high structural similarity with surface-layer homology (SLH) domain trimers found in bacterial Surface (S)-layer proteins (Supplementary Figure 6) (Kern et al., 2011; Sychantha et al., 2018). Bacterial Surface (S)-layer proteins are known to have glycan binding capabilities, therefore, based on the structural similarity, the helical domain of GUS may also have role in substrate recognition or positioning.

Though, there is no systematic study, there are few reports indicating role of C-terminal domain in the substrate processing and oligomeric assembly. The catalytic activity of Rg3GUS (PDB ID: 6MVG) was drastically reduced upon deletion of DUF4982 (Ervin, Li, et al., 2019). The CBM57 domain in BdGUS (PDB ID: 6ED1) contributes a loop that extends into the active site, potentially participating in the substrate recognition and processing (Biernat et al., 2019). DUF2 in PmGUS and CBM57 in BdGUS (PDB ID: 6ED1) have been shown to participate in the oligomeric organization (Little et al., 2018; Biernat et al., 2019). Overall, this analysis indicates that while the GH2 core remains conserved, the C-terminal extensions are the primary source of structural diversity and functional specialization among microbial GUS enzymes. It is important to note that the functional role of many of these C-terminal domains is not completely understood and requires further biochemical and structural characterization. Still, this new domain architecture-based classification provides a useful framework for understanding GUS diversity. Earlier, evolutionary diversity of β-galactosidases has been suggested to be driven by the addition of different non-catalytic domains at the C-terminal region (Talens-Perales et al., 2016). A similar model of C-terminal domains-directed evolution can be envisaged for β-glucuronidases.

### The diversity of active site loops

As mentioned earlier, the GUS active site is situated in the cleft of the TIM-barrel domain and is adorned by several loops, typically 8-25 residues in length. These loops are proven to be one of the critical determinants of substrate specificity and processivity in different GUS enzymes (Pollet et al., 2017). These β/ɑ -type loops originate primarily from the TIM barrel domain. However, in some recent studies, loops originating from N-terminal ancillary domains have also been shown to contribute to the substrate binding (Pellock et al., 2018). Walker et. al., (2022) classified gut microbial GUS enzymes into eight categories primarily based on the presence and length of active site loop arrangements (Table 2 and Supplementary Figure 7). These categories are labeled as Loop1 (L1), mini Loop1 (mL1), Loop2 (L2), mini Loop2 (mL2), mini Loop1,2 (mL1,2), No Loop (NL), N-terminal loop (NTL) and No Loop FMN (NL-FMN). Based on this classification, we assigned different loops to the GUS enzymes identified in the present study using a multiple sequence alignment approach and validated through comparative structural analyses of computational models (Figure 4).

**Figure 4:**
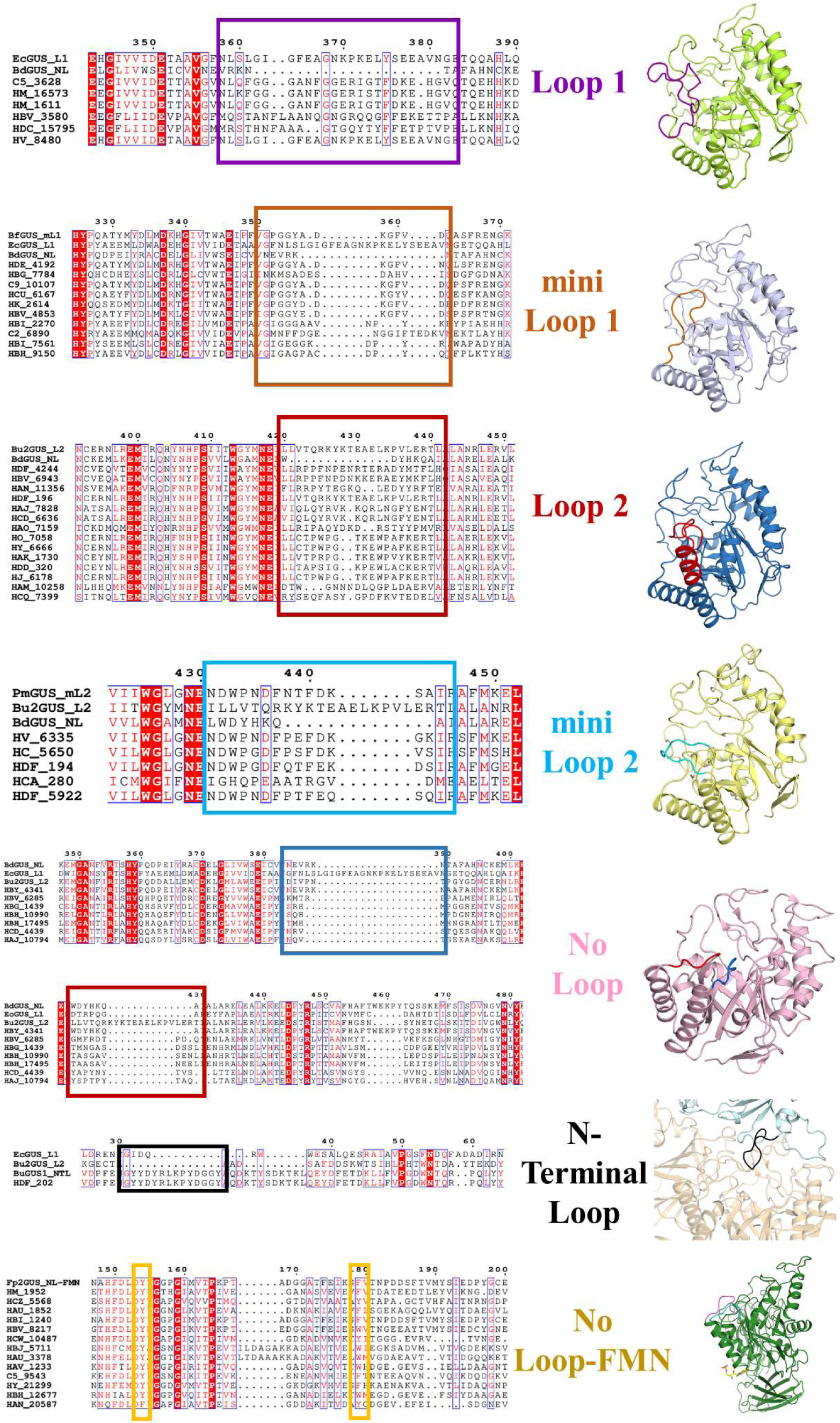
Characterization of different loop types among GUS enzymes identified in the current study using multiple sequence alignment with representative sequence of each loop type. Loop regions are highlighted by colored boxes over MSA and also by different color in the structural model of selected homologs of each loop type. For clarity, only TIM-barrel domain is shown.

**Table 2:**
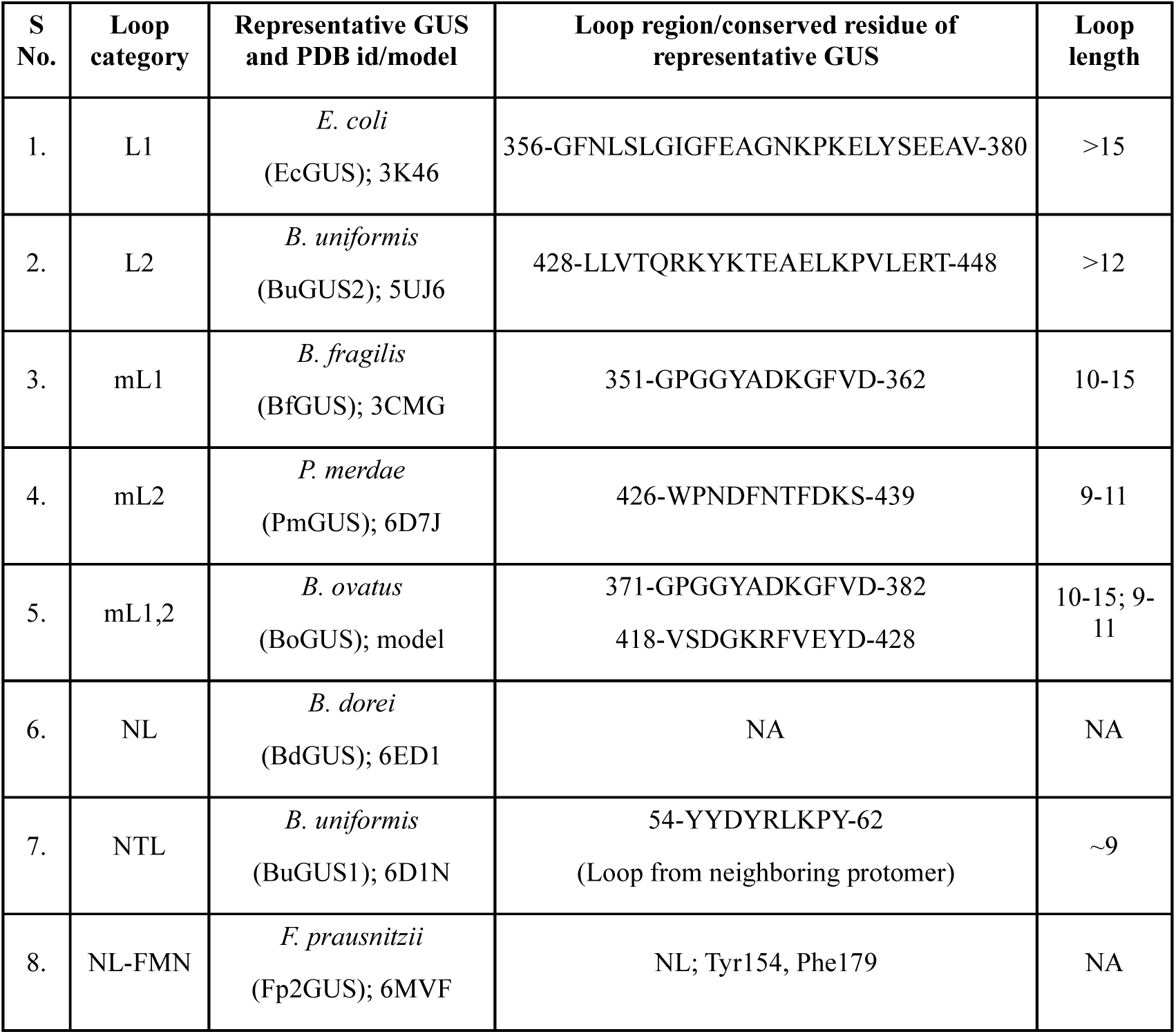
Loop categories and their representative GUS proteins.

The analyses revealed a diverse spectrum of loop architectures, associated with specific taxonomic distributions and functional implications. L1-GUS enzymes, predominantly found in proteobacteria (e.g., *Escherichia coli*) and firmicutes (e.g., *Faecalibacterium, Eubacteria, Streptococcus*), maintain consistent loop lengths of more than 15 residues but vary in amino acid composition. These loop variations likely facilitate the recognition of diverse small-molecule glucuronides, such as drug metabolites. Mini-Loop 1 (mL1) variants are more abundant comprising 10 sequences (12%) of the total than the L1-GUS (6 sequences, 7%). L2-GUS enzymes, representing 18% of the dataset (14 sequences), are prevalent in Bacteroidetes and Actinobacteria (e.g., *Prevotella, Bacteroides* and *Bifidobacterium*) while mL2 enzymes account for 6% (5 sequences) and are primarily identified in *Bacteroides* and *Akkermansia*. The most abundant category is NL-GUS, representing 33 sequences (41%). Ten sequences belong to the subset of NL-FMN GUS while only one sequence was identified that falls in the NTL loop category. Recently, in our lab, we have identified the presence of a unique loop in *Akkermansia muciniphila* termed as the J-loop as it originates from the jelly-roll domain (Supplementary figure 8) (Tarushi et al., 2025). This loop is unique to *Akkermansia* and not present in any other known GUS enzymes. AmGUS was able to cleave drug-glucuronide *i.e.,* SN-38G efficiently and the J-loop was suggested to assist in this processing. Discovery of this new loop suggests yet unexplored structural and functional diversity in microbial GUS enzymes.

The distinct distribution of these loop architectures across different bacterial phyla reflects population-specific microbial compositions, likely shaped by host-specific factors such as diet, lifestyle, and genetics. The high prevalence of the NL and L2 architectures suggests an evolutionary adaptation for processing larger, more complex glycoconjugates compared to the small-molecule glucuronide processed primarily by L1-GUS. Further biochemical characterization is required to establish the precise correlation between loop-type abundance, taxonomic distribution, and dietary glycan preferences.

### Distribution of FMN-NL GUS enzymes

A specialized subset of NL-GUS is FMN-binding GUS which exhibits unique substrate preferences, including the ability to cleave glucuronide derivatives of anticancer drug regorafenib, immunosuppressant mycophenolate mofetil (MMF) and antibacterial triclosan (Ervin, Hanley, et al., 2019; Xia et al., 2024; Zhang et al., 2022). In these enzymes, flavin mononucleotide (FMN) binds to a surface groove involving conserved aromatic residues, e.g., Tyr154 and Phe179 in Fp2GUS (PDB ID: 6MVF) (Pellock et al., 2019). Twenty-three potential FMN-binding GUS enzymes were identified based on the presence of these conserved aromatic residues using a MSA based approach. Further, structural analysis showed that the FMN binding site is a shallow surface groove and should be freely accessible (Tarushi et al., 2025). Therefore, we have used the presence of an accessible surface groove as an additional criterion to identify ‘true’ NL-FMN GUS. This approach identified 10 putative NL-FMN GUS harboring the conserved aromatic residues and accessible surface groove (Figure 4, Supplementary figure 9). For all other GUS enzymes, FMN-binding motifs are either not conserved and/or the groove is absent/inaccessible due to variations in adjacent structural elements (Supplementary figure 9). For example, in AmGUS, despite having high homology with NL-FMN GUS enzymes, the surface groove is inaccessible due to the presence of a longer β-strand coming from the Ig-like domain (Tarushi et al., 2025). This structure-guided bioinformatics identification of FMN-binding GUS enzymes provides a foundation for future experimental validation by site-directed mutagenesis and biochemical activity assays.

### Identification of calcium-binding GUS enzymes

Beyond the GUS rubric, active site loops and FMN-binding site, certain GUS enzymes including Bu2GUS (*Bacteroides uniformis*, L2, PDB ID: 5UJ6) harbor a unique calcium-binding site located approximately 24 Å from the active site (Pellock et al., 2018). This metal-binding site has been shown to be essential for catalytic activity. The site-directed mutagenesis and structural studies revealed that the calcium-binding site is defined by three aspartate residues and three water molecules. For the identification of Ca^2+^ binding sites within 79 GUS enzymes, we performed MSA using Bu2GUS as a reference. The sequences were probed for the presence of the coordinating triad: Asp176, Asp341, and Asp367 (numbering per Bu2GUS) and further verified by structural analysis (Figure 5A). Our analysis identified 10 GUS enzymes, all belonging to the L2-GUS category, that possess the conserved residues required for Ca^2+^ coordination (Figure 5B). The exact mechanism by which this Ca^2+^ binding influences the GUS activity, however, is not known. The Ca^2+^-binding sites, however, is consistently associated with the L2 loop-type architecture. Interestingly, all the Ca^2+^-binding L2-GUS enzymes identified in this study also contain a CBM57 domain at their C-terminus. The concurrent presence of the L2 loop, CBM57 domain, and Ca^2+^-binding sites suggests a specialized evolutionary adaptation within the Bacteroidota phylum where these structural features appear to have co-emerged as an integrated functional module. Further investigations are needed to understand the combined role of these elements in modulating GUS activity and the evolutionary pressures that have driven their co-emergence.

**Figure 5:**
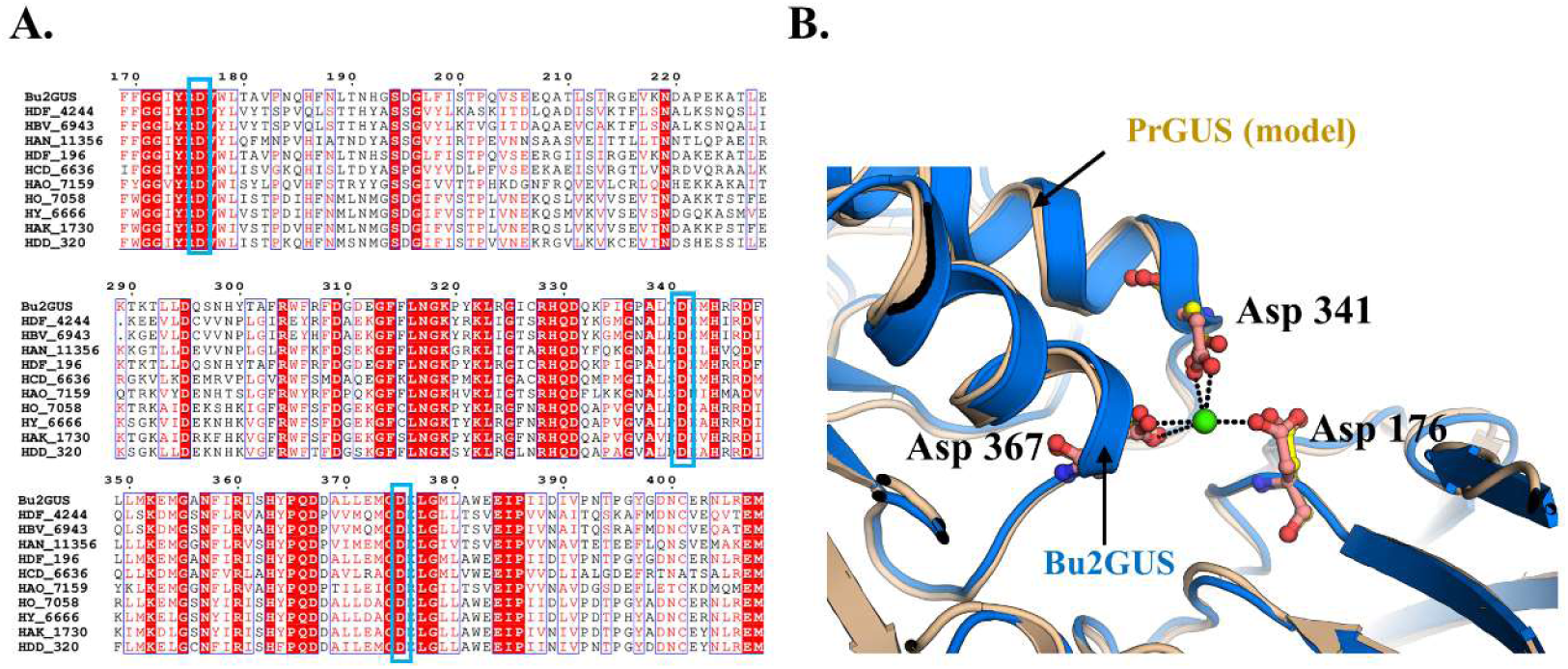
Identification of Ca^2+^-binding sites in the putative GUS enzymes. (A) Multiple sequence alignment of GUS enzymes with Bu2GUS (PDB ID: 5UJ6) showing conservation of aspartic acid triad (corresponding to residues Asp176, Asp341, and Asp367). (B) Ca^2+^-binding site in BuGUS2 (*blue*). Three conserved aspartates (Asp176, Asp341, and Asp367) along with three waters coordinating Ca^2+^ are shown. Predicted Ca^2+^-binding site of *Prevotella* GUS enzyme (PrGUS, HAO_7159 (model), *wheatish*) is also shown.

### Sequence similarity network

To gain an understanding of the sequence-structure-function relationship among the identified GUS enzymes, the sequence similarity network (SSN) was created using the EFI-EST tool (Oberg et al., 2023; Zallot et al., 2019). An alignment score of 10^-130^ separated the 79 GUS enzymes into six clusters (Figure 6). Interestingly, these six clusters broadly correspond to the classification based on the domain architecture proposed in the current study. The class 1 GUS enzymes, which contain only the GH2 core were bifurcated into two clusters. One cluster consisted solely of NL-GUS, while the other clusters includes both L1- and mL1-GUS enzymes. Similarly, class 2 GUS with DUF1 domain were also divided into two clusters; one composed entirely of NL-GUS while other included predominantly NL-GUS along with a few mL1-GUS enzymes. The Class 3 GUS containing the CBM57 domain formed an independent cluster belonging almost exclusively to the L2-GUS category, with an exception of a single sequence that is identical to BdGUS and classified as NL-GUS. Interestingly, BdGUS also possess CBM57 at the C-terminal ends which is a characteristic of L2-GUS enzymes. The class 4 GUS having the DUF2 domain formed a separate cluster comprising solely of the mL2-loop category of GUS. The class 5 GUS, characterized by the presence of helical domains were all NL-GUS and clustered together with a subset of Class 2 NL-GUS enzymes.

**Figure 6:**
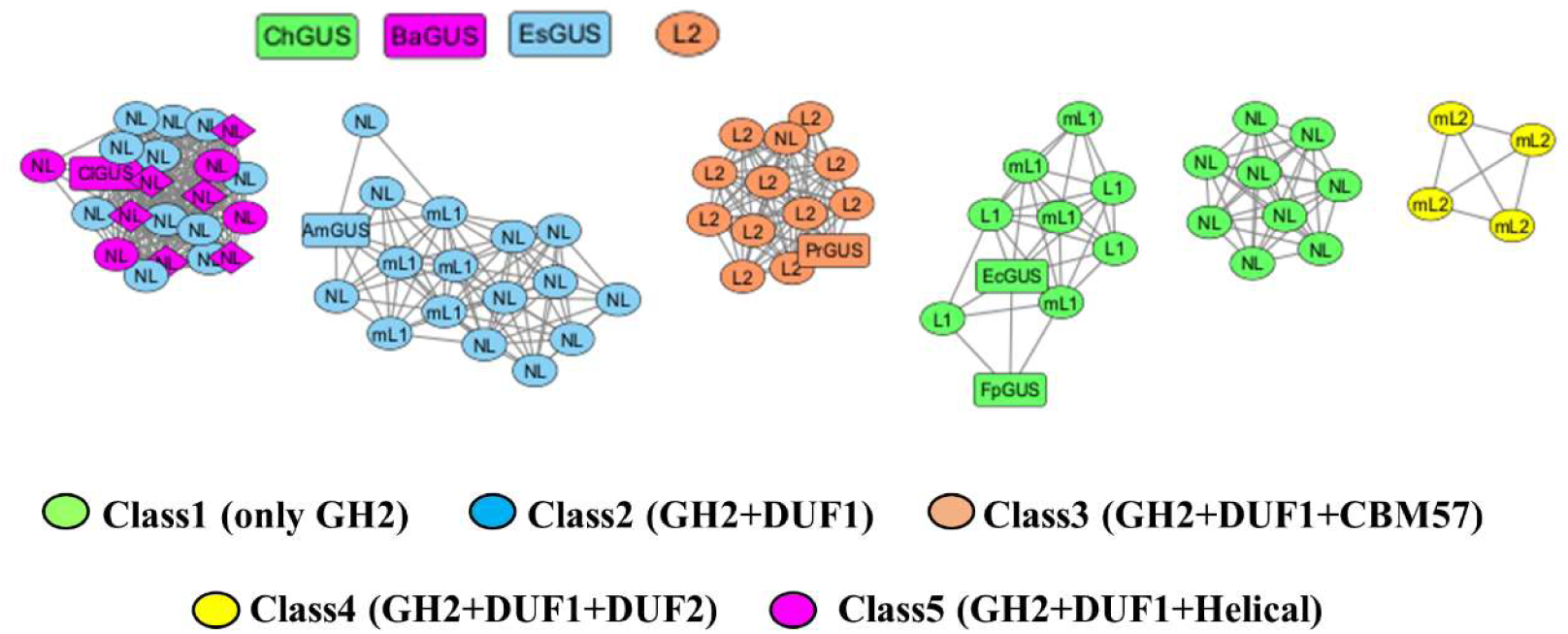
Sequence similarity network analysis of 79 microbial GUS enzymes showing six major clusters. The color represents different classes based on the domain architectures. Each GUS is identified by the loop type mentioned in the circle. NL-FMN GUS are represented in diamond shape. The GUS highlighted in rectangles have been selected for in vitro characterization discussed in later section.

Overall, the SSN analysis suggests a strong association between domain architecture and loop-type among GUS enzymes. NL-GUS exhibited the greatest structural diversity, encompassing all domain architecture classes except Class 4, whereas the mL1 GUS showed only Class 1 and Class 2 type domain architectures. Other loop categories displayed a much stronger association with specific domain architectures. Presence of CBM57 domain was strongly correlated with the L2 loop while the DUF2 domain was present only in the mL2-GUS enzymes. These patterns suggest that domain architecture and loop configuration may have co-evolved during GUS diversification with specific domain-loop combinations likely conferring distinct substrate recognition properties. Further phylogenetic and functional studies are required to validate evolutionary relationship between these structural features and to determine their relative contribution in providing substrate diversity to GUS enzymes.

### Conserved structural framework beyond GUS rubric

While the conservation of “GUS rubric” residues is well-characterized, our multiple sequence alignment of 79 diverse GUS enzymes revealed a suite of strictly conserved residues outside the canonical framework. Based on their spatial arrangements and hydrogen-bonding networks, these residues can be categorized into two functional groups: active-site proximal residues and inter-domain stabilizing residues (Supplementary Table 1). Active site proximal residues either interact directly with the substrate or are positioned adjacent to the “GUS rubric” and likely maintain active-site geometry through a secondary shell of interactions. Among them, His335 interacts directly with the substrate and its orientation is stabilized by hydrogen bonding with Glu356. (Figure 7A; residue numbering corresponding to RgGUS, PDB ID: 6JZ5). His301, on the other hand, forms water-mediated hydrogen bonds with the ligand and its orientation is stabilized by hydrogen bonding with Tyr336. Substrate recognition is further supported by interaction with Asp167, which also participates in a hydrogen-bonding network with Arg570 and Asn574 (the N*KG motif), (Figure 7B). Arg332 stabilizes the orientation of GUS rubric residues Asn417 and Glu512 and forms a network involving Glu356 and water mediated interactions with Thr511 and Gly299 (Figure 7C). These conserved interactions are likely to influence the proper positioning of ligand-binding residues and facilitate optimal substrate binding. Further, His301, Arg332, and His335 may also provide a necessary positive electrostatic environment to bind the highly electronegative, oxygen-rich glucuronide moiety of the substrate.

**Figure 7:**
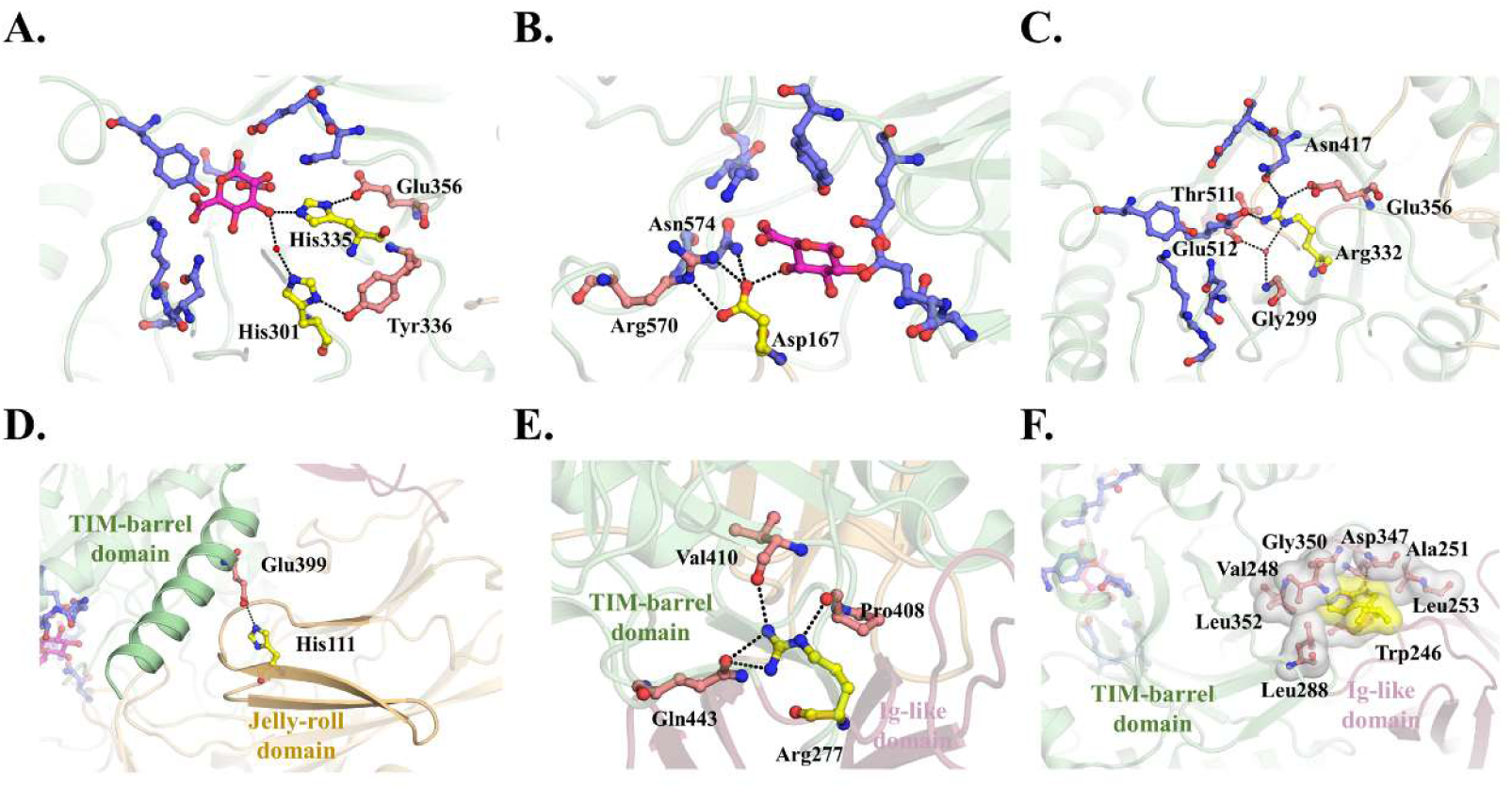
Conservation analysis of residues beyond GUS rubric. (A) Interactions of His301 and His335 with the substrate. (B) Interactions of Asp167 with the substrate. (C) Stabilization of GUS rubric residues by Arg332. (D) His111 from Jelly-roll domain interacts with Glu399 of the TIM-barrel domain. (E) Interactions of Arg277 from Ig-like domain with residues Pro408, Val410, and Gln443 of the TIM-barrel domain. (F) Trp246 from Ig-like domain is sequestered by a hydrophobic patch created by non-polar residues from the TIM-barrel domain.

Beyond substrate binding, several conserved residues appear to participate in inter-domain stabilization. Present at the interfaces of the jelly-roll, Ig-like, and TIM-barrel domains, these residues form interactions required for maintaining the relative orientation of different domains. For instance, His111, located on a β-strand of the jelly-roll domain forms a stabilizing hydrogen bond with Glu399 from the TIM-barrel domain (Figure 7D). Similarly, Arg176, extending from another β-strand in the jelly-roll domain, anchors the N-terminus to the catalytic core through a network of direct and water-mediated hydrogen bonds involving Glu302, Pro337, Phe115, Glu340, Glu341, and Arg403. Similar interactions are observed at the interface between the Ig-like domain and the TIM-barrel domain. Tyr254, located at the base of a β-strand from the Ig-like domain, interacts with Arg244, possibly stabilizing the loop, which interacts with the TIM-barrel domain. Similarly, Arg277 from the Ig-like domain interacts with the Gln443, Pro408 and Val410 of the TIM-barrel domain (Figure 7E). Furthermore, Trp246 of Ig-like domain is sequestered within a conserved hydrophobic patch formed by Val248, Leu253, Leu288, Gly350, Leu352 and Pro408 from the TIM-barrel and forms hydrogen bonding with Asp347 (Figure 7F). While Gly350 is absolutely conserved, the surrounding residues are conservatively replaced by other hydrophobic amino acids, suggesting structural importance of this hydrophobic packing. A few other residues including some glycines, are also found to be conserved (Supplementary Table 1). Though not immediately apparent from the current structural analysis, their conservation suggests an important role in the structural integrity of GUS enzymes. Therefore, though the lack of experimental data prevents definitive functional assignments, these structural insights provide a robust framework for future studies to establish their specific functional contributions.

### Cellular localization and physicochemical properties

To understand the cellular localization of different GUS enzymes, the presence of N-terminal signal peptide was predicted using the SignalP 6.0 server (Teufel et al., 2022). No L1 enzymes were found to possess a signal sequence, implying that L1 proteins are retained within the intracellular environment. In contrast, the majority of L2, and mL2 GUS exhibited signal sequences, indicating the potential for translocation across the inner microbial membrane. These findings suggest a functional divergence, wherein periplasmic proteins may be specialized for acting on larger polysaccharide substrates, while intracellular enzymes likely target smaller glucuronides capable of cellular entry (Supplementary Figure 10). Distribution of key physicochemical properties like isoelectric point (pI), aliphatic index and GRAVY values of GUS enzymes indicate towards soluble, mesophilic enzymes specifically adapted to the human gastrointestinal environment (Supplementary Figure 11, Supplementary Table 2). Analysis of isoelectric point (pI) distribution showed a predominantly acidic profile, with 68% of sequences falling between 5.0 and 6.5, aligning with the mildly acidic pH of the gut lumen. The enzymes exhibit consistent mesophilic thermostability, indicated by aliphatic indices between 63.39-87.37 and universal hydrophilicity, as evidenced by exclusively negative GRAVY scores ranging from −0.256 to −0.665. The wide variation in the molecular weights from ∼62 to ∼102 kDa—reflect diverse domain architectures including canonical 60–70 kDa GH2 unit and additional C-terminal modules.

### Expression and purification of putative GUS enzymes

Based on the domain architecture, loop-type and low sequence similarity with the known GUS structures, eight putative GUS enzymes were selected for the detailed in vitro characterization studies (Supplementary Table 3). For biochemical characterization, heterologous expression and purification of GUS enzymes were performed in *E. coli* BL21 (DE3) cells. Expression of GUS enzymes was explored at two different temperatures, i.e., 37℃ and 18℃. The recombinant proteins were purified using nickel affinity chromatography and the results were evaluated through SDS-PAGE. Seven out of eight GUS proteins were expressed as soluble proteins at both temperatures; however, at 37℃, most of the expressed proteins were in the insoluble fraction except for AmGUS (Supplementary figure 12). For all the GUS enzymes, the yield of soluble proteins was higher at 18°C (Figure 8A, Supplementary Figure 12). The GUS proteins were further purified by size exclusion chromatography and stored in storage buffer (50 mM Hepes, 100 mM NaCl, pH 7.5). The storage buffers used for EcGUS and AmGUS were 50 mM Pipes, 100 mM NaCl, pH 6.5 and 50 mM Mes, 100 mM NaCl, pH 6.0 respectively. The expression of ChGUS and EsGUS need further optimization as level of expression for ChGUS was much lower, even at 18°C and no expression was observed for EsGUS.

**Figure 8:**
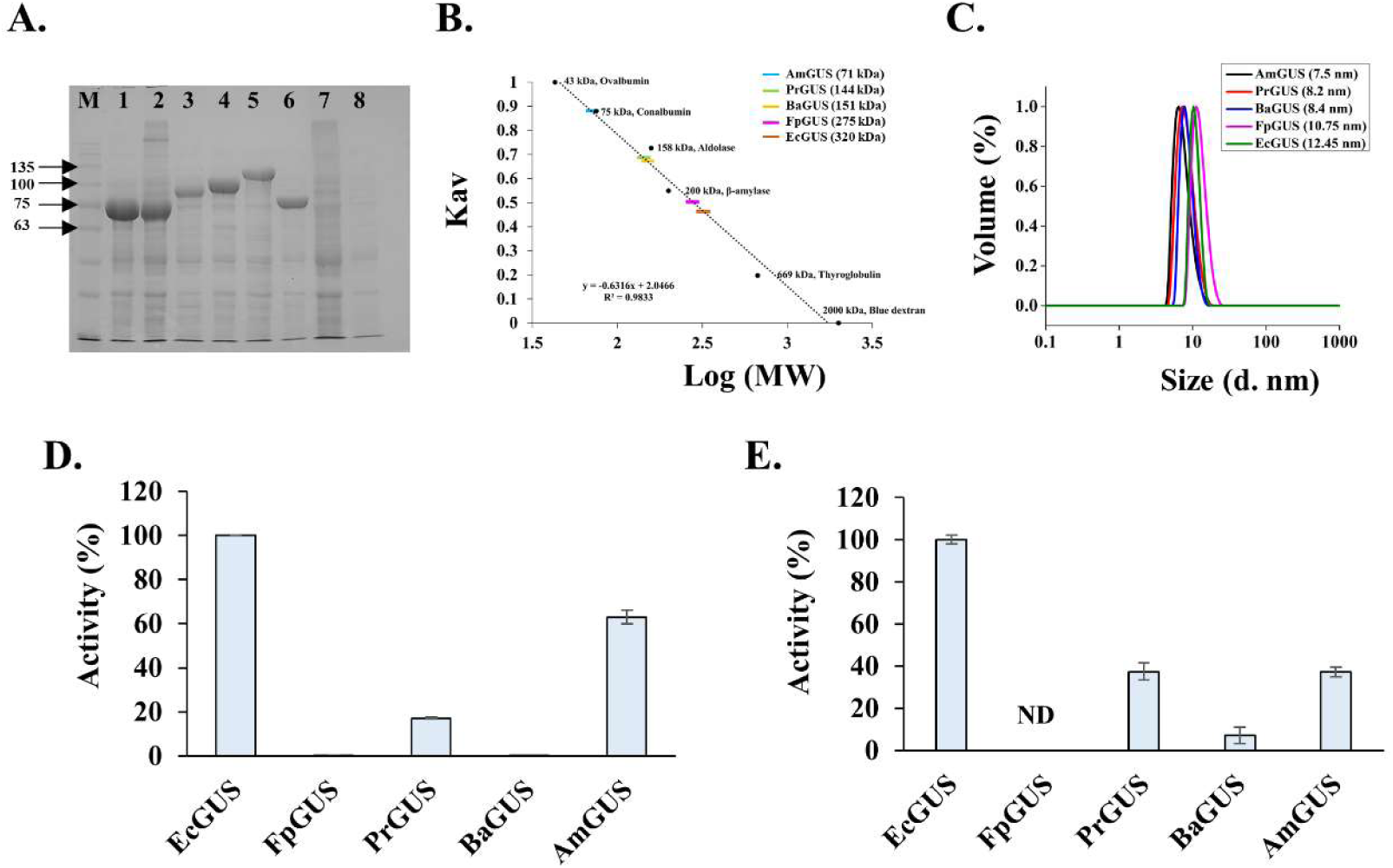
In vitro characterization of selected GUS panel. (A) SDS-PAGE showing soluble fractions of expressed GUS enzymes. M – Marker; 1 – EcGUS, ∼71 kDa; 2 – FpGUS3, ∼72 kDa; 3 – ClGUS, ∼88 kDa; 4 – PrGUS, ∼99 kDa; 5 – BaGUS, ∼107 kDa; 6 – AmGUS, ∼78 kDa; 7 – ChGUS, ∼65 kDa; 8 –EsGUS, ∼83 kDa (B) Determination of the oligomeric nature of the identified GUS enzyme using SEC calibration curve. The molecular weights estimated using calibration curve are given in the brackets. (C) DLS profiles of different GUS enzymes showing homogeneous protein preparations. The estimated hydrodynamic diameters are mentioned in the brackets. (D) In vitro GUS activity of purified GUS enzymes with 4-MUG. (E) In vitro SN38-G processing activity of purified GUS enzymes. Experiments were done in triplicates at their respective optimum pH. Error bars represent mean ± SD. Biological replicates were also conducted.

The oligomerization characteristics of the purified proteins were evaluated using size-exclusion chromatography and dynamic light scattering, revealing that GUS enzymes exist in multiple oligomeric states (Figure 8B and 8C). GUS enzymes have been shown to adapt different oligomeric states including dimer, trimer, tetramer, and hexamer (Wang et al., 2021; Pellock et al., 2019). In our study, FpGUS3 along with EcGUS showed tetrameric state while PrGUS and BaGUS may have monomeric or compact dimeric organization as SEC and DLS results are inconclusive. Recently, we have reported that AmGUS show potential monomeric form, first for a GUS enzyme (Tarushi et al., 2025). Oligomeric nature of ChGUS and ClGUS could not be characterized due to poor expression and multiple peaks in gel filtration respectively. The oligomeric states of these GUS enzymes need to be further verified by other methods including multi-angle light scattering (MALS) or ultra-centrifugation.

### Validation of GUS activity

The representative panel of seven GUS enzymes were examined for their activity using fluorogenic substrate, 4-methylumbelliferone-glucuronide (4-MUG, SRL India). All seven proteins showed positive activity with 4-MUG confirming that the putative GUS enzymes are indeed active β-glucuronidases. ClGUS and ChGUS also showed GUS activity, however, they were not further characterized due to purification and stability issues. Activity optima were determined for five GUS enzymes within the pH range of 4.0–10.0 using CHC buffer. The GUS enzymes showed activity across a broad range of pH 5.0-8.5 with optimum pH varying in a narrow range of 5.5-6.5 (Supplementary figure 13). The optimum pH range may be related to the localization of microbial species along the gastrointestinal tract with pH ranging from ∼5.0 to 7.4. Relative activities of five purified GUS enzymes were evaluated at their respective optimum pH using 4-MUG as substrate. In comparison to EcGUS, AmGUS and PrGUS showed low to moderate levels of activity while FpGUS3 and BaGUS showed very low activity (Figure 8D).

### SN-38G cleavage activity

The active metabolite of the anti-cancer drug irinotecan (CPT-11) is SN-38, which is glucuronidated to SN-38G, during the detoxification process in the liver. Reactivation of this SN-38G into SN-38 by gut microbial GUS activity has been demonstrated to be a major factor in the irinotecan-induced intestinal toxicity. The SN-38G processing ability of our GUS panel is determined by an optimized fluorescence-based assay. In addition to EcGUS, AmGUS and PrGUS showed significant SN-38G hydrolysis while BaGUS showed poor activity (Figure 8E, Table 3). FpGUS3, despite having an L1 loop and showing high homology with EcGUS, showed no activity. As reported previously, AmGUS belongs to the mL2 category and its SN-38G processing is attributed to the presence of a unique J-loop over the active site (Tarushi et al., 2025). PrGUS belongs to the L2 loop category and is the second L2-GUS enzyme characterized to have SN-38G activity (Bhatt et al., 2020). Identification of two new GUS enzymes with SN-38G processing activity broadens the GUS diversity capable of processing drug-glucuronides. However, crystal structures of AmGUS and PrGUS are required to establish molecular determinants of SN-38G processing activity of AmGUS and PrGUS.

**Table 3:** Characteristics of purified GUS enzymes.

| **GUS enzyme** | **Organism** | **Domain class** | **Loop category** | **Optimum pH** | **SN-38G activity** |
| --- | --- | --- | --- | --- | --- |
| EcGUS | *Escherichia coli* | GH2 | L1 | 6.5 | +++++ |
| FpGUS | *Faecalibacterium prausnitzii* | GH2 | L1 | 6.0 | NA |
| PrGUS | *Prevotella* | GH2+DUF1+CBM57 | L2 | 6.5 | ++ |
| BaGUS | *Bifidobacterium adolescentis* | GH2+DUF1+helical domain | L2 | 5.5 | NA |
| AmGUS | *Akkermansia muciniphila* | GH2+DUF1 | mL2 | 6.5 | ++ |

## Conclusion

In this study, we identified a set of new GUS enzymes from a representative metagenomic database of the Indian population. Domain architecture analysis using computational structural modelling revealed presence of different types of C-terminal domains. A new classification for microbial GUS enzymes based on the presence and nature of additional non-catalytic domains at the C-terminal end has been proposed and mapped with existing loop-based categorization. The cellular localization as well as physicochemical properties have been predicted. The in vitro characterization of selected putative GUSs was explored through recombinant expression in *E. coli.* Seven GUS enzymes showed activity on 4-MUG validating the bioinformatics approach for identification of ‘true’ GUS enzymes. The GUS panel showed optimum activity in a narrow pH range of 5.5 to 6.5. Further, two GUS enzymes other than EcGUS showed drug-glucuronide (SN-38G) processing activity. We have identified several conserved residues other than the ‘GUS rubric and provided structural basis for their possible role in substrate recognition and inter-domain stabilization. Identification of new domain architecture and loop-type have expanded the potential structural functional diversity of GUSome in the human microbiome and provide the basis for their functional characterization. Moreover, a possible evolutionary and functional link between domain architecture and loop-type is suggested. Further, biochemical and biophysical characterization will help in validation of these observations and establish a complete structure-function paradigm of GUS enzymes.

## Materials and methods

### Identification of GUS enzymes

For the identification of GUS enzymes, a representative gene catalogue of the Indian gut microbiome was chosen. In a cohort study of 110 individuals, Dhakan and his colleagues have established a gene catalogue of the Indian gut microbiome representing the uniqueness of the Indian population in comparison to other populations (Dhakan et al., 2019). This gene catalogue was downloaded from GigaDB (https://gigadb.org/) and non-redundant protein sequences were then used to create a local database for BLASTp searches. For the identification of GUS homologs, 24 unique known GUS sequences belonging to 17 different bacteria were used as query sequences. In the first step, GUS homologs were selected using an E-value cutoff of <0.05 and a sequence identity threshold of ≥25%. In the next step, the obtained GUS homologs were then clustered to a 95% sequence identity threshold using CD-HIT (v3.2.1) and used for downstream analysis (Fu et al., 2012; Li & Godzik, 2006). Each protein sequence was then checked for the presence of the N*KG motif in a pairwise alignment with one of the query GUS sequences using NCBI-BLASTp (Altschul et al., 1990). The other GUS rubric residues were then checked in a multiple sequence alignment using Clustal Omega (Sievers et al., 2011) via the EMBL-EBI server (Madeira et al., 2019) along with the 24 query sequences. Computational structural models of identified GUS enzymes were prepared with AlphaFold2 as implemented in ColabFold and using an in-house deployment with a default set of parameters (Jumper et al., 2021; Mirdita et al., 2022). For sequences with more than 99% identity to known GUS enzymes, crystal structures available in the Protein Data Bank (Berman et al., 2000) were used directly for analysis. For other sequences, models in the AlphaFold database (Varadi et al., 2022) with more than 99% identity were used directly. Comparative structural analysis with known GUS structures was performed in PyMol 3.2.0 and WinCOOT to verify the presence of properly organized active site including GUS rubric (Emsley et al., 2010). We termed protein sequences that satisfied the above conditions as ’true’ β-glucuronidases.

### Taxonomic Assignments of GUS sequences

Taxonomy for identified GUS enzymes was assigned using NCBI-BLASTp search against the non-redundant (nr) database. We examined the top results showing identity of more than 95% to define the genera and 99% to determine the bacterial species. However, if sequences with <95% identity were found, the genus was not assigned and is denoted as undefined. The analysis is based on the nr database as of May 2026 (Altschul et al., 1990).

### Domain categorization

The identified GUS sequences were analyzed for domain architecture using a combined sequence-structure based workflow. Initially, protein sequences were subjected to the dbCAN3 server for automated CAZyme annotation (Zheng et al., 2023) dbCAN3 assigns different domains based on Hidden Markov Model (HMM) searches against the CAZy database. The default set of parameters were used for dbCAN3 server. To further validate and improve the annotation confidence, the sequences were analyzed using InterProScan (Blum et al., 2025). To examine the spatial arrangement and domain boundaries identified from dbCAN3 and InterProScan, the structural models were visualized in PyMol 3.2.0 (Schrödinger, LLC, 2025). Structural inspection enabled confirmation of the canonical GH2 core domain along with other C-terminal domains where applicable. Further, the server “Foldseek” was used to identify domain similar to the helical domains observed in identified GUS enzymes (van Kempen et al., 2024). This integrated computational workflow was employed to group the GUS proteins into separate architectural categories.

### Loop classification

For loop classification, the identified GUS sequences were aligned with the representative GUS enzymes categorized according to loop-types (Table 2). Second-step verification was done in a pairwise alignment approach against the 279 catalogued and annotated GUS enzymes from the HMP stool sample database (Pollet et al., 2017). We considered alignment criteria with an E-value < 0.05 and the highest percentage identity to be valid hits against the 279 catalogued sequences for the loop classification. The sequence-basis loop classified proteins were further validated on a structural basis using structural models and comparing with the reference structures. Structural analysis was performed with WinCOOT and PyMol 3.2.0.

### Identification of NL-FMN GUS

For the identification of NL-FMN GUS enzymes, GUS homolog sequences were screened for the presence of the conserved FMN-binding residues, *i.e.,* residues aligning with Tyr154 and Phe179 of *Faecalibacterium prausnitzii* (Fp2GUS) in an MSA using Clustal Omega. Presence of any of the aromatic residues, *i.e.,* tyrosine, phenylalanine or tryptophan was accepted. A second step of filtering was performed by verifying the presence of an accessible surface groove in computational models. We classified the GUS homologs with the conserved aromatic residues and accessible surface groove at the FMN binding site as the NL-FMN GUS.

### Prediction of signal peptide and physicochemical properties

The sequences were assessed for the presence of signal peptide cleavage sites using the SignalP 6.0 online server (Teufel et al., 2022). Various physicochemical properties like theoretical molecular weights, pI and aliphatic index were also deduced using SignalP 6.0, while GRAVY score was determined using the online Protein GRAVY tool (https://stothardresearch.ca/sequence-manipulation-suite/#tool=protein-hydropathy).

### Expression and purification of selected GUS enzymes

All the putative GUS sequences selected for the in vitro studies were codon-optimized, synthesized and subcloned into the expression vector (pET21b) with an N-terminal hexahistidine tag and TEV protease cleavage site using Nde1 and Xho1 restriction sites. The protein sequences were analyzed for the signal peptide using SignalP 6.0 and omitted while cloning. The recombinant plasmids carrying specific GUS gene inserts were transformed into *E. coli* BL21 (DE3) cells. Primary cultures (10 ml) of these transformed *E. coli* cells were grown overnight in Luria-Bertani (LB) broth supplemented with ampicillin (100 μg/ml) at 37°C with shaking at 100 rpm. Following day, a small inoculum (1:1000 v/v) was used to inoculate secondary culture. The cells were grown at 37°C in orbital shaker at 120 rpm. At OD600=0.8, the temperature was reduced to 18°C and after 30 minutes, the cells were supplemented with isopropyl-β-D-1 thiogalactopyranoside (IPTG) to a final concentration of 0.5 mM for protein induction and continued for overnight growth at 18°C. The next morning, cells were pelleted via centrifugation at 12000xg and stored at −40°C until further use. For protein purification, the cell pellets were resuspended in lysis buffer (50 mM Tris, 200 mM NaCl, pH 7.5) and disrupted via sonication. The cleared whole-cell lysate was obtained at 4°C by centrifugation at 12000xg and loaded onto the Ni-NTA column and binding was allowed for 30 min at 4°C on the rocker. The column was given two washes using wash buffer (50 mM Tris, 200 mM NaCl, 20 mM imidazole, pH 7.5). The proteins were eluted using elution buffer (50 mM Tris, 200 mM NaCl, 350 mM imidazole, pH 7.5). The purity was checked on 10% SDS-PAGE and the eluted fractions were pooled, concentrated and loaded on the Biorad ENrich SEC 650 column connected to the Akta purifier 10 system. The proteins were eluted at a flow rate of 0.5 ml per minute and their molecular weights were estimated using a calibration curve.

### Dynamic Light Scattering

Dynamic light scattering (DLS) experiments were carried out on the Zetasizer Nano-ZS instrument (Malvern Pananalytical Ltd., UK) in which the backscattering angle of the scattered light was set at 173°. The purified GUS proteins were centrifuged at 30000 x g for 30 min before the data collection. The concentration of 1 mg/ml of each GUS was used and the experiment was performed in a low-volume disposable sizing cuvette. Initially, an equilibration time of 30 sec was given followed by two measurements at 10°C (with every measurement having 14 runs). The size distribution graph representing the relative volume (%) against the diameter (nm) was plotted.

### GUS activity using 4-MUG assay

The GUS activity was measured using fluorescence substrate, 4-MUG. The experiments were performed in 96-well, flat-bottomed, black fluorescence plates. The total reaction volume was 50 µl in 50 mM CHC buffer (Citric acid: HEPES: CHES – 2:3:4 molar ratio) and 50 mM NaCl with the final concentration of the enzyme specific to each GUS. The reaction was kept constant at room temperature and initiated by the addition of 0.5 mM 4-MUG. The reaction was stopped after 10 min using 2 M sodium carbonate and the fluorescence signal was measured (λ_Ex_: 360nm; λ_Em_: 450nm) in a Clariostar multimode reader (BMG Labtech, Germany). Experiments were done in triplicates and repeated twice.

### SN-38G processing assay

The SN-38G processing ability is determined by an optimized fluorescence-based assay in 96-well, flat-bottomed, black plates. The total reaction volume was 50 µl in 50 mM CHC buffer and 50 mM NaCl with the final concentration of the 1nM of each GUS. The experiment was performed at room temperature and the reaction was initiated by the addition of 50 µM SN-38G. The reaction was stopped after 10 min using 2 M sodium carbonate and the fluorescence was measured (λ_Ex_: 360 nm; λ_Em_: 420 nm) in a Clariostar multimode reader (BMG Labtech, Germany). Experiments were done in triplicates and repeated twice.

## Contributions

**Tarushi:** Writing – original draft, Methodology, Investigation, Formal analysis, Visualization. **Chinmaya V Badgujar:** Investigation**. Subhash C Bihani:** Conceptualization; Formal analysis; Investigation; Methodology; Visualization; Writing – original draft; review & editing.

## Acknowledgments

We are thankful to Dr Mukesh Kumar, and Dr Gagan Deep Gupta, PCS, Biosciences Group, Bhabha Atomic Research Centre (BARC), for insightful discussions. We gratefully acknowledge the Computer Division, BARC for providing computational resources for the bioinformatics study. The work is funded by the Department of Atomic Energy, India.

