## Supplementary data for "Expanding the GUSome: Structure-guided identification and characterization of gut microbial β-glucuronidases"

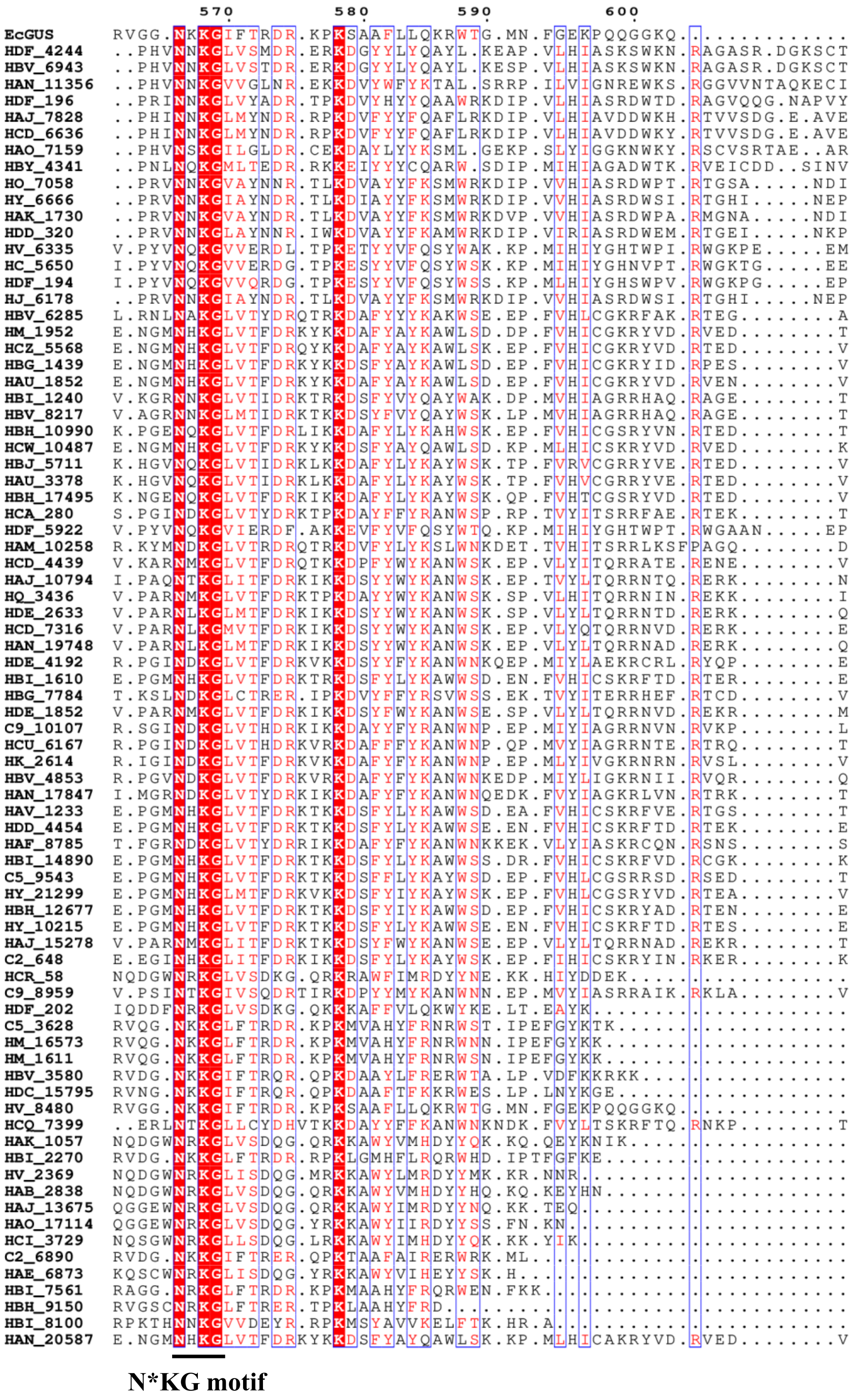


Supplementary Figure 1:
Multiple sequence analysis showing the conservation of N*KG motif among the identified 79 GUS homologs

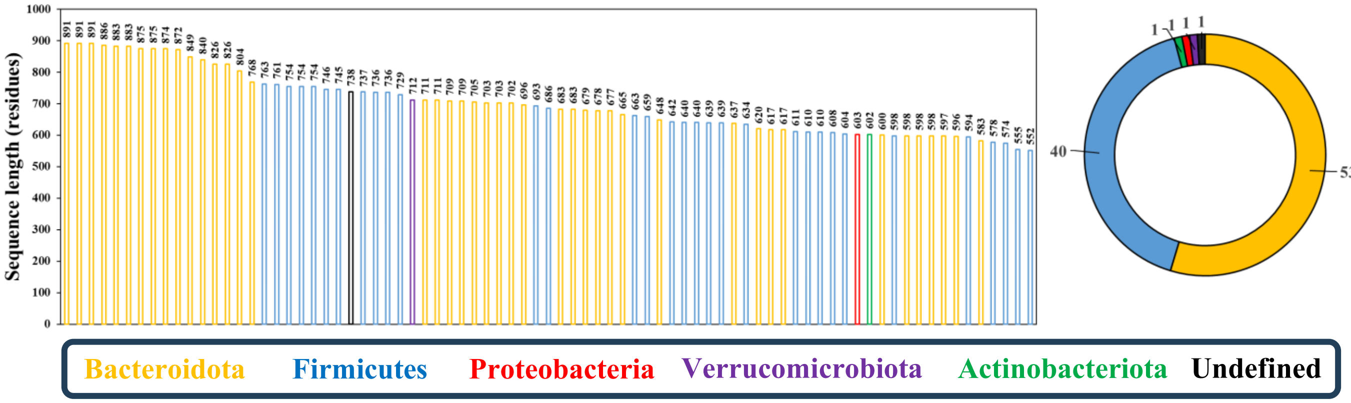


Supplementary Figure 2:
Taxonomic classification of identified 79 GUS sequences. Sequences are arragned according to the length of the proteins. Sequences belonging to different phyla are represented in different colors. Values indicated in the pie chart correspond to the percentage distribution.

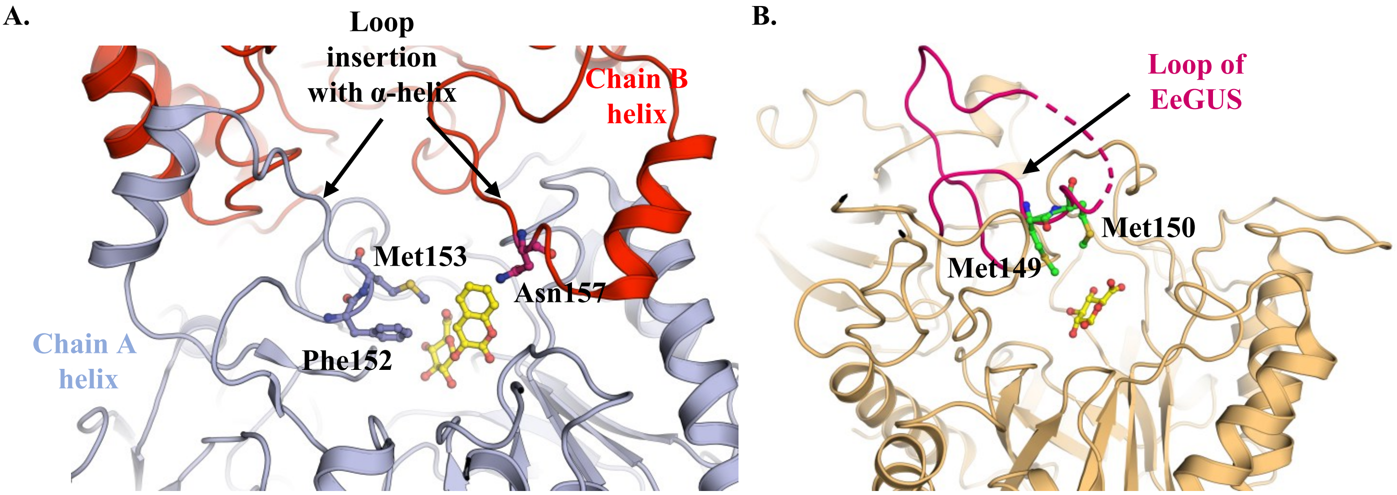


Supplementary Figure 3:
Role of loop insertion of jelly-roll domain in substrate binding: (A) The helical loop insertion in the jelly-roll domain of BiGUS (*Bifidobacterium dentium,* PDB ID: 6LD6, light blue and red). Residues from the loop insertion of the same chain (*light blue*) and from neighboring chain (magenta) enter into the active site and interact with the bound ligand (*yellow*). (B) The loop insertion in EeGUS (*Eubacterium eligens,* PDB ID: 6BJW, *magenta*), the residues from the insertion (*green*) are oriented towards the active site and likely to interact with the bound ligand (*yellow*).

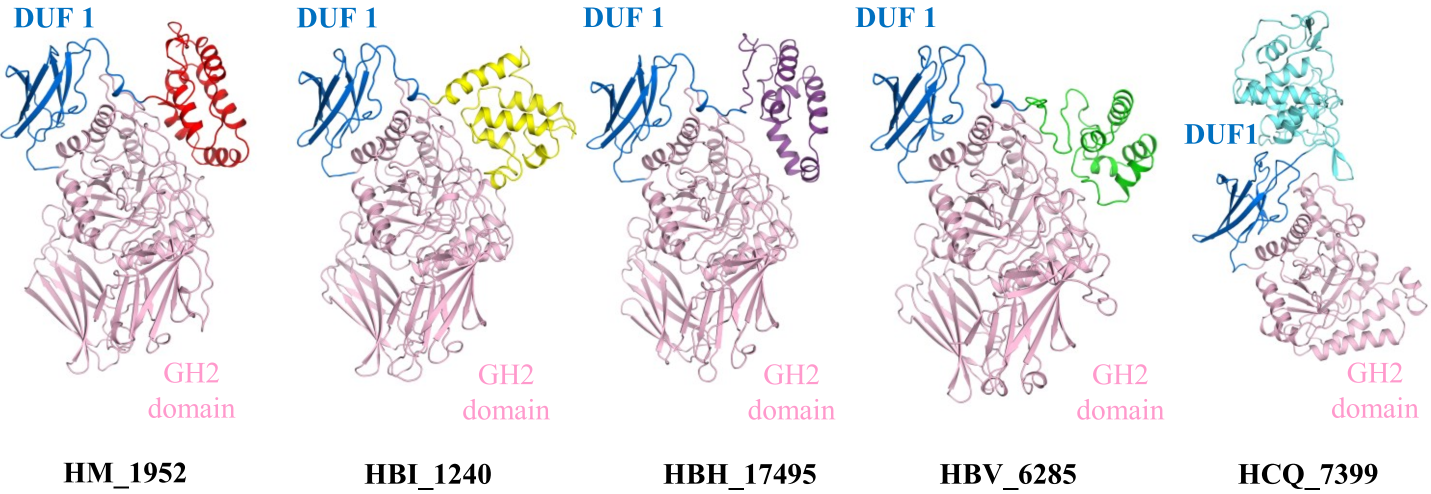


Supplementary Figure 4:
Different type of C-terminal helical domains identified in the present study. Figures are the cartoon representation of computational models. GH2 domain (*pink*); DUF1 (*blue*), Helical domain is shown in different colors.

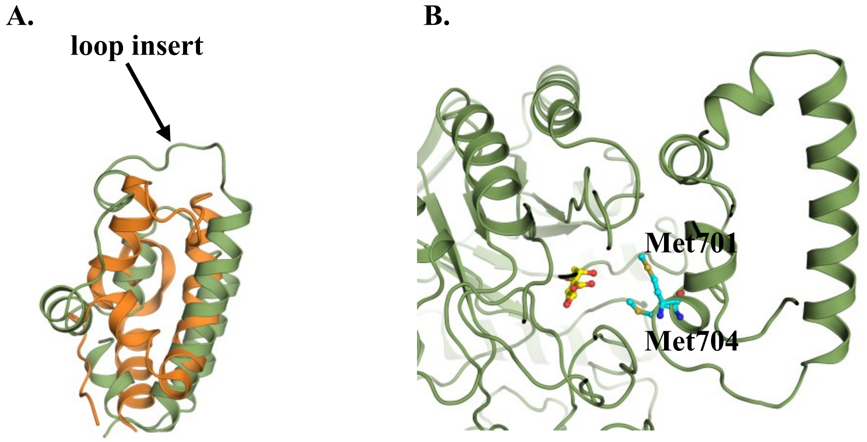


Supplementary Figure 5:
C-terminal helical domain of Class 5 GUS enzymes: (A) Superposition of C-terminal helical domain of HBG_1439 (model, *smudge green*) with C-terminal domain of GH3 β-glucosidase from *Clostridium thermocellum* (PDB ID: 7MS2, *orange*) highlighting similarity between the two domains. (B) Two methionine residues (*cyan*) from the loop insert of helical domain in HBG_1439 are positioned suitably to interact with the aglycone moiety of substrate suggesting role of C-terminal helical domain in the substrate binding.

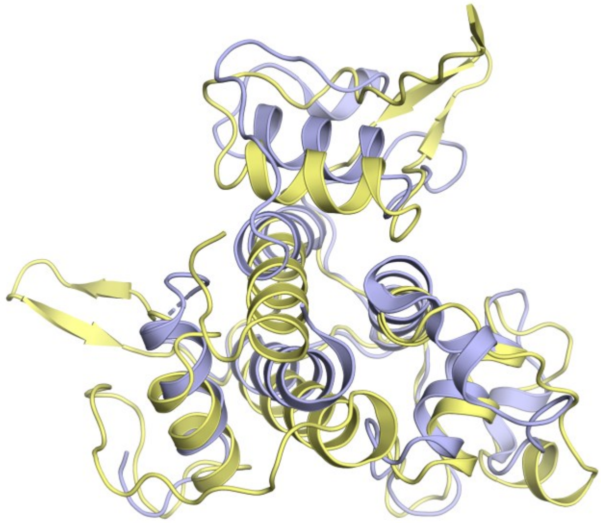

Supplementary Figure 6:
C-terminal helical domain of *Bifidobacterium adolescentis* GUS (HCQ_7399 (model), *pale yellow*) show striking similarity with surface-layer homology (SLH) domain trimers of bacterial surface (S) layer proteins (PDB ID: 6BT4, *light blue*).

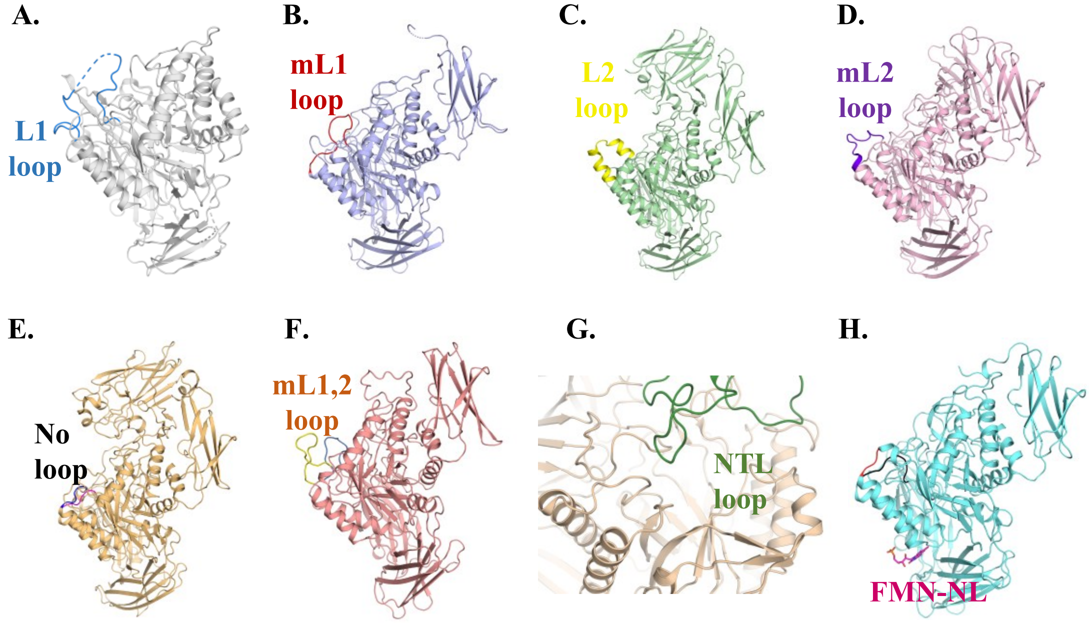


Supplementary Figure 7:
Representative GUS enzymes of different loop types. (A) L1-GUS (EcGUS, PDB ID: 6LEM); (B) mL1-GUS (BfGUS, PDB ID: 3CMG); (C) L2-GUS (Bu2GUS, PDB ID: 5UJ6); (D) mL2-GUS (PmGUS, PDB ID: 6D7J); (E) NL-GUS (BdGUS, PDB ID: 6ED1); (F) mL1,2-GUS (BoGUS, model); (G) NTL-GUS (BuGUS1 PDB ID: 6D1N); (H) FMN-NL-GUS (Fp2GUS, PDB ID: 6MVF)

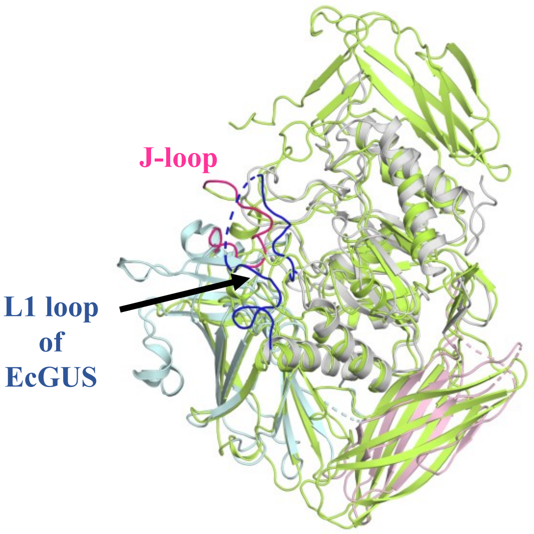


Supplementary Figure 8:
J-loop of AmGUS originates from jelly-roll domain, occupies similar topological space as that of L1 loop of EcGUS and is likely to interact with the aglycone moiety of the substrate.

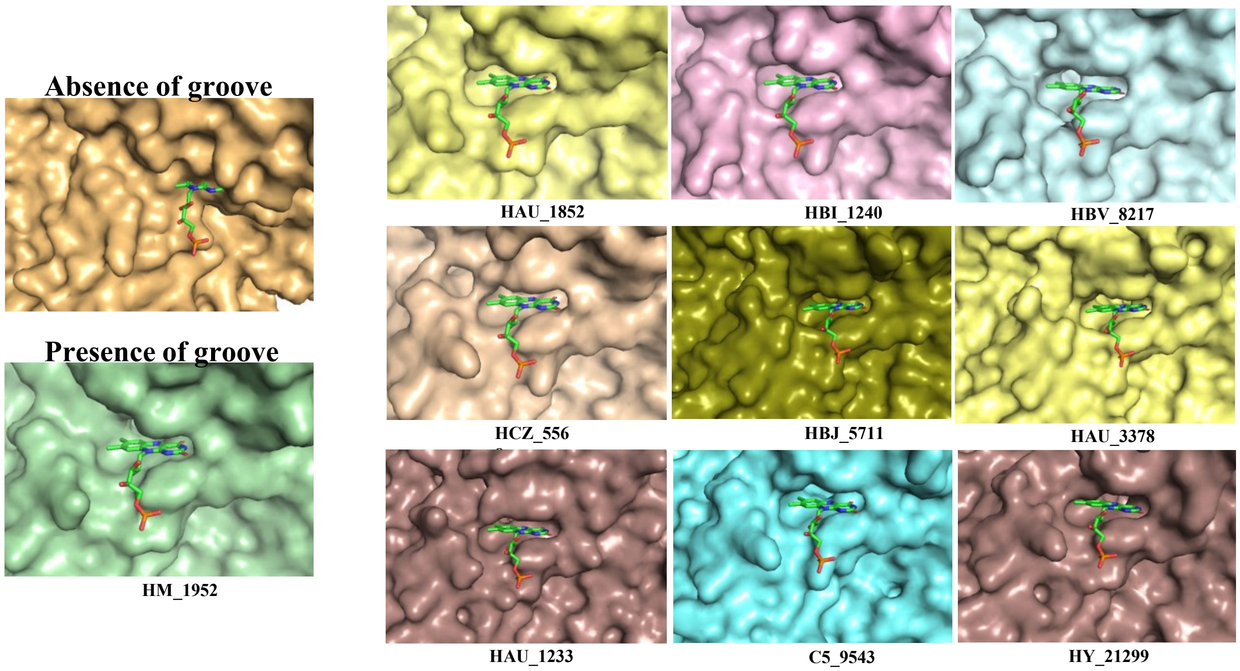


Supplementary Figure 9:
Surface representation of a FMN-binding NL-GUS (HM_1952; model) highlighting the surface groove and a non-FMN binding GUS (*Clostridium* GUS; model) showing absence of groove. FMN is docked by structural superposition with (PDB ID: 6MVF). Remaining FMN-binding NL-GUS identified in the present study are also shown.

| Sr No. | Residue (RgGUS) | Residue (EcGUS) |
|---|---|---|
| 1 | His111 | His107 |
| 2 | Gly113 | Gly109 |
| 3 | Asp167 | Asp163 |
| 4 | Gly173 | Gly169 |
| 5 | Arg176 | Arg172 |
| 6 | Trp246 | Trp242 |
| 7 | Tyr254 | Tyr250 |
| 8 | Arg277 | Arg272 |
| 9 | Asn289 | Asn284 |
| 10 | His301 | His296 |
| 11 | Arg332 | Arg327 |
| 12 | His335 | His330 |
| 13 | Gly350 | Gly345 |
| 14 | Glu356 | Glu351 |
| 15 | Trp413 | Trp408 |
| 16 | Asp441 | Asp436 |
| 17 | Trp477 | Trp471 |
| 18 | Tyr478 | Tyr472 |
| 19 | Gly514 | Gly506 |
| 20 | Trp557 | Trp549 |
| 21 | Asp561 | Asp553 |
| 22 | Arg570 | Arg562 |
| 23 | Lys586 | Lys578 |

Supplementary Table 1:
Conserved residues beyond GUS rubric.

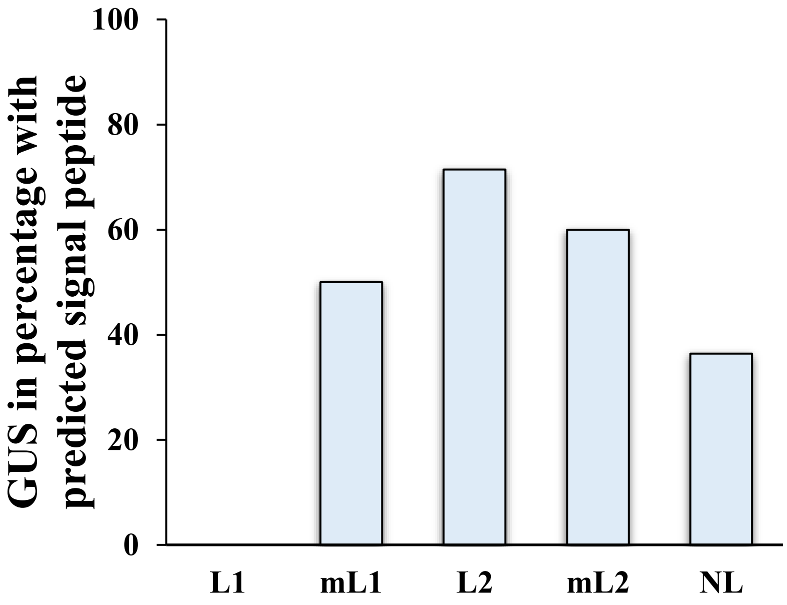


Supplementary Figure 10:
Signal peptide analysis of identified GUS enzymes

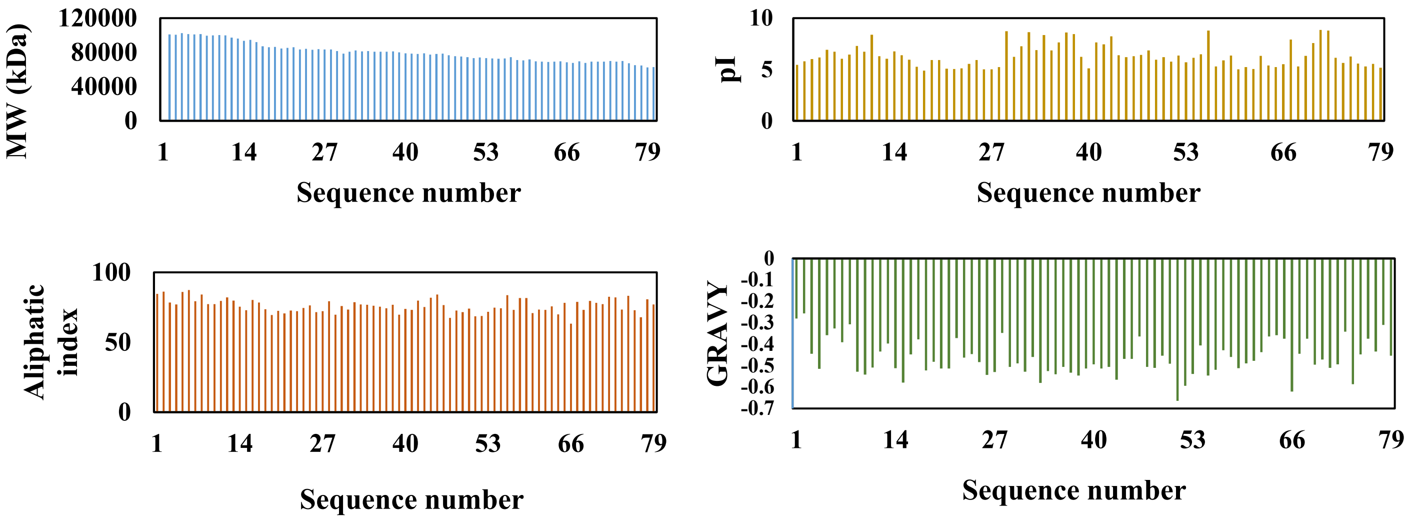


Supplementary Figure 11:
Physicochemical properties of identified GUS proteins

| Property | Range Observed | Key Interpretation |
|---|---|---|
| Molecular Weight | 62-102 kDa | Multi-domain diversity; correlates with domain architecture |
| Isoelectric Point (pI) | 4.8 – 8.84 | Predominantly acidic; adapted to gut lumen pH |
| Aliphatic Index | 63.39-87.37 | Moderately thermostable; mesophilic character |
| GRAVY Score | −0.256 to −0.665 | Universally hydrophilic; soluble |

Supplementary Table 2:
Physicochemical properties of identified GUS enzymes

| S. No. | Gene ID | Length (in residues) | Organism | Loop-type | PDB homolog (% identity) |
|---|---|---|---|---|---|
| 1. | HV_8480 | 625 | *Escherichia coli* (EcGUS) | L1 | 6LEG (100%) |
| 2. | HDC_15795 | 626 | *Faecalibacterium prausnitzii* (FpGUS3) | L1 | 6EC6 (68%) |
| 3. | HBG_1439 | 776 | *Clostridia* (ClGUS) | NL | 6MVG (59%) |
| 4. | HAO_7159 | 876 | *Prevotella* (PrGUS) | L2 | 6D50 (34%) |
| 5. | HCQ_7399 | 965 | *Bifidobacterium adolescentis* (BaGUS) | L2 | 3CMG (34%) |
| 6. | HCA_280 | 710 | *Akkermansia muciniphila* (AmGUS) | mL2 | 3CMG (40%) |
| 7. | HBI_8100 | 577 | *Christensenellaceae* (ChGUS) | NL | 6NCW (47%) |
| 8. | HBG_7784 | 722 | *Eubacteria* (EsGUS) | mL1 | 3CMG (34%) |

Supplementary Table 3:
A representative set of GUS sequences selected for in vitro characterization.

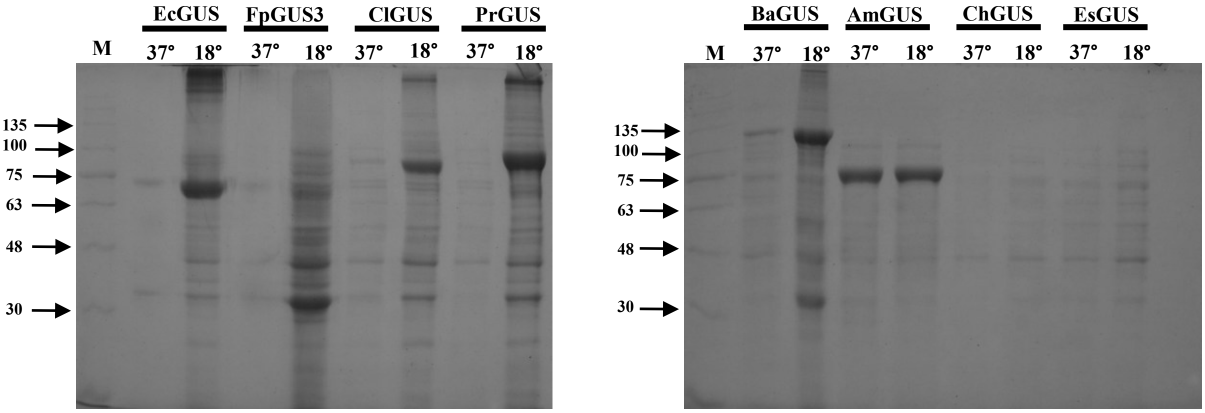


Supplementary Figure 12:
10% SDS–PAGE differential expression levels of the recombinant proteins at two different temperatures, 37°C and 18°C. Only the solube fractions obtained after cell lysis and centrifugation is loaded for each protein. M-protein ladder.

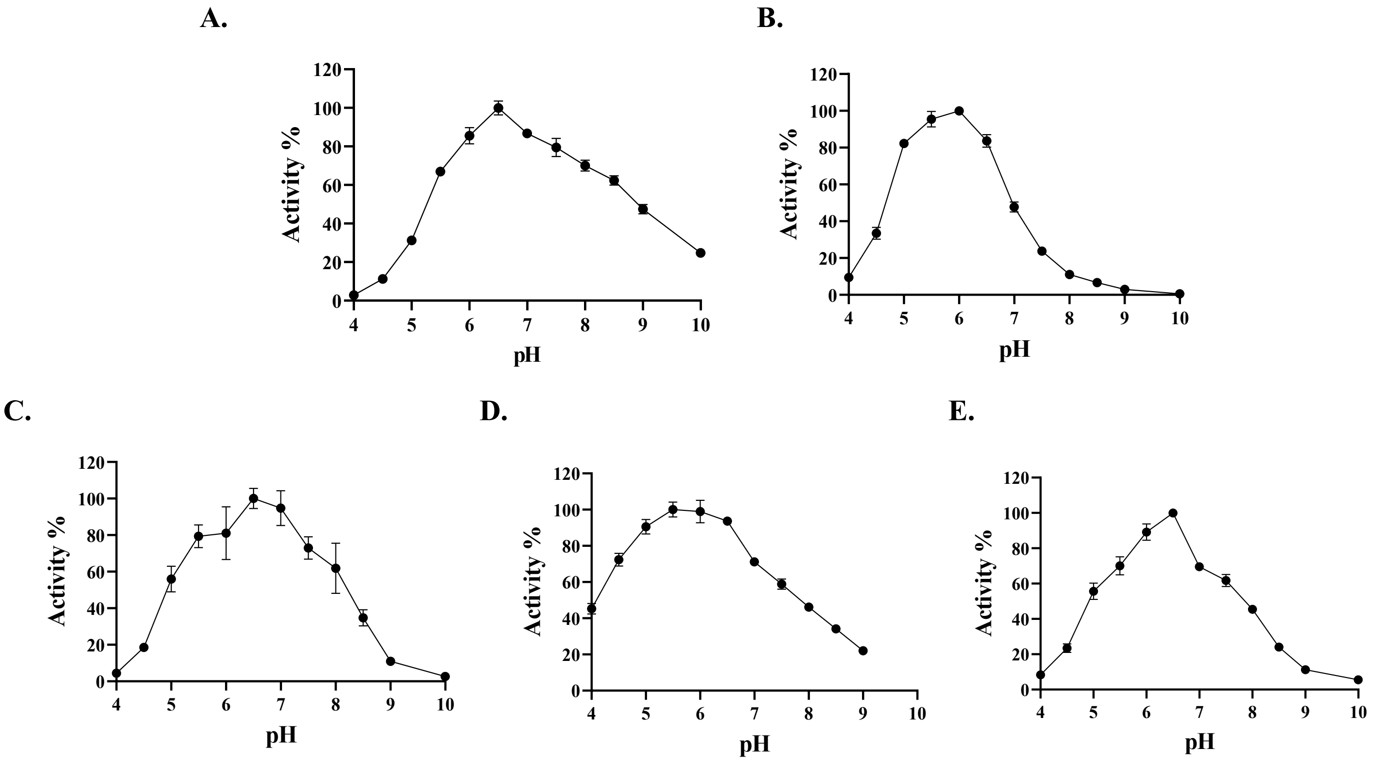


| **GUS** | EcGUS | FpGUS3 | PrGUS | BaGUS | AmGUS |
|---|---|---|---|---|---|
| **pH** | 6.5 | 6.0 | 6.5 | 5.5 | 6.5 |

Supplementary Figure 13:
Determination of the optimum pH of the identified GUS enzymes. (A) EcGUS; (B) FpGUS3; (C) PrGUS; (D) BaGUS; (E) AmGUS. Experiments were done in triplicate and error bars represent mean ± SD.
